# Human immunodeficiency virus-1 induces host genomic R-loop and preferentially integrates its genome near the R-loop regions

**DOI:** 10.1101/2024.03.06.583715

**Authors:** Kiwon Park, Dohoon Lee, Jiseok Jeong, Sungwon Lee, Sun Kim, Kwangseog Ahn

**Author notes:** Corresponding author (KA). These authors contributed equally to this work (Kiwon Park, Dohoon Lee).

## Abstract

Although HIV-1 integration sites favor active transcription units in the human genome, high-resolution analysis of individual HIV-1 integration sites has shown that the virus can integrate into a variety of host genomic locations, including non-genic regions. The invisible infection by HIV-1 integrating into non-genic regions, challenging the traditional understanding of HIV-1 integration site selection, is more problematic because they are selected for preservation in the host genome during prolonged antiretroviral therapies. Here, we showed that HIV-1 integrates its viral genome into the vicinity of R-loops, a genomic structure composed of DNA– RNA hybrids. VSV-G-pseudotyped HIV-1 infection initiates the formation of R-loops in both genic and non-genic regions of the host genome and preferentially integrates into R-loop-rich regions. Using a HeLa cell model that can independently control transcriptional activity and R-loop formation, we demonstrated that the exogenous formation of R-loops directs HIV-1 integration-targeting sites. We also found that HIV-1 integrase proteins physically bind to the host genomic R-loops. These findings provide novel insights into the mechanisms underlying retroviral integration and the new strategies for antiretroviral therapy against HIV-1 latent infection.

## Introduction

Retroviruses cause permanent host infections by integrating their reverse-transcribed viral genomes into the host genome. Retroviral integration considerably affects a wide range of biological phenomena, including the persistence of fatal human diseases and the shaping of metazoan evolution (*1*). Human immunodeficiency virus (HIV)-1 is a representative retrovirus underlying the global burden of acquired immune deficiency syndrome (AIDS) (*2*). The chromosomal landscape of HIV-1 integration has been shown to influence proviral gene expression, the persistence of integrated proviruses, and prognosis in antiretroviral therapy (*3–5*). Integration into the host genome is not random and displays distinct preferences for gene-dense regions, where active transcription occurs (*6*), by interacting with host factors, such as transcription activators, epigenetic marker binding proteins, and super enhancers (*7–13*). However, transcriptional activity is not the sole determinant of the HIV-1 integration site landscape (*10*). For instance, some highly expressed genes do not necessarily correspond to a high level of integration in PBMCs infected in culture with HIV (*14*). Despite their lower probability of integration, HIV-1 proviruses are observed in non-genic regions of the genomes of infected individuals (*4*). This indicates the possibility of there being an undiscovered mechanism or determinant that constitutes the correct genomic environment for HIV-1 integration.

An R-loop is a three-stranded nucleic acid structure that comprises a DNA–RNA hybrid and a displaced strand of DNA, and has long been considered a transcription byproduct (*15, 16*). R-loops in cellular genomes are enriched in actively transcribed genes as they occur naturally during transcription (*15, 17*). However, R-loop formation is not limited to gene body regions and is widespread in the genome (*15*). As a result of *in trans* R-loop formation, R-loops are also abundant in non-genic regions, such as intergenic regions and repetitive sequences, including transposable elements, centromeres, and telomeres (*15, 18–20*), independent of the transcriptional activity of the genes harboring the R-loops. Although R-loops have been identified as critical intermediates and regulators of a number of biological processes (*15, 16, 21*), the dynamics and the role played by cellular R-loops in pathological contexts remain unclear.

R-loops are important contributors to molding the genomic environment and spatial organization of the cellular genome, and can potentially play a novel role in host-pathogen interaction. In the cellular genome, R-loops relieve superhelical stresses and are often associated with open chromatin marks and active enhancers (*22, 23*), which are also distributed over HIV-1 integration sites (*6, 9, 10*). In transcription-induced R-loop formation, a guanine-quadruplex (G4) structure can be generated in the non-template DNA strand of the R-loop (*24*). A recent study showed that G4 DNA can influence both productive and latent HIV-1 integration (*25*). In addition, R-loops are prevalent non-canonical B-form DNA structures (*26*) and intermediates between the B-form DNA and A-form RNA conformations (*27*). It has recently been reported that the B-to-A transition in the target DNA occur during retroviral integration (*27, 28*). Accumulating evidence suggests that host genomic R-loops are undiscovered host factors in the HIV-1 integration site selection mechanism that dynamically interact with the host genomic environment.

Here, we showed a notable role for R-loops in the interaction between HIV-1 and its host, specifically in HIV-1 integration. VSV-G-pseudotyped HIV-1-infection induces host cellular R-loop formation, and the R-loop rich regions of the host genome are preferred for HIV-1 integration in diverse cell types. HIV-1 integrase proteins showed considerable binding affinity for nucleic acid substrates containing R-loop structures. Our results suggest that R-loops are important components of the host genomic environment for HIV-1 integration site determination.

## Results

### Host genomic R-loops accumulate by VSV-G-pseudotyped HIV-infection in diverse cell types

To investigate the relationship between HIV-1 infection and host cellular R-loops, we first analyzed R-loop dynamics in different types of cells infected with HIV-1 at early post-infection time points using DNA–RNA immunoprecipitation followed by cDNA conversion coupled with high-throughput sequencing (DRIPc-seq) using a DNA–RNA hybrid-specific binding antibody, anti-S9.6 (*29*). HeLa cells, primary CD4^+^ T cells isolated from two individual donors, and the CD4^+^/CD8^−^ T cell lymphoma Jurkat cell line were infected with the VSV-G-pseudotyped HIV-1-EGFP in a time-course manner. The infected cells were harvested at the same post-seeding time point but at 0, 3, 6, and 12 h post infection (hpi) for DRIPc-seq library construction (Fig. 1A and S1A-C Fig.). Our DRIPc-seq analysis yielded loci-specific R-loop signals at the referenced R-loop-positive loci (RPL13A and CALM3) and an R-loop-negative locus (SNRPN) (*29*), which were both strand-specific and highly sensitive to pre-immunoprecipitation *in vitro* RNase H treatment in HeLa cells, CD4^+^, and Jurkat T cells (Table S1-3). Notably, the number of DRIPc-seq peaks mapped to the human reference genome increased markedly during the early post infection with HIV-1 (3 and 6 hpi for HeLa cells and 6 and 12 hpi for CD4^+^ and Jurkat T cells; Fig. 1B). Most of the peaks that mapped in cells harvested at 0 hpi were commonly found in all other samples, but a significant numbers of unique peaks were observed after infection (Fig. 1C).

**Fig 1.**
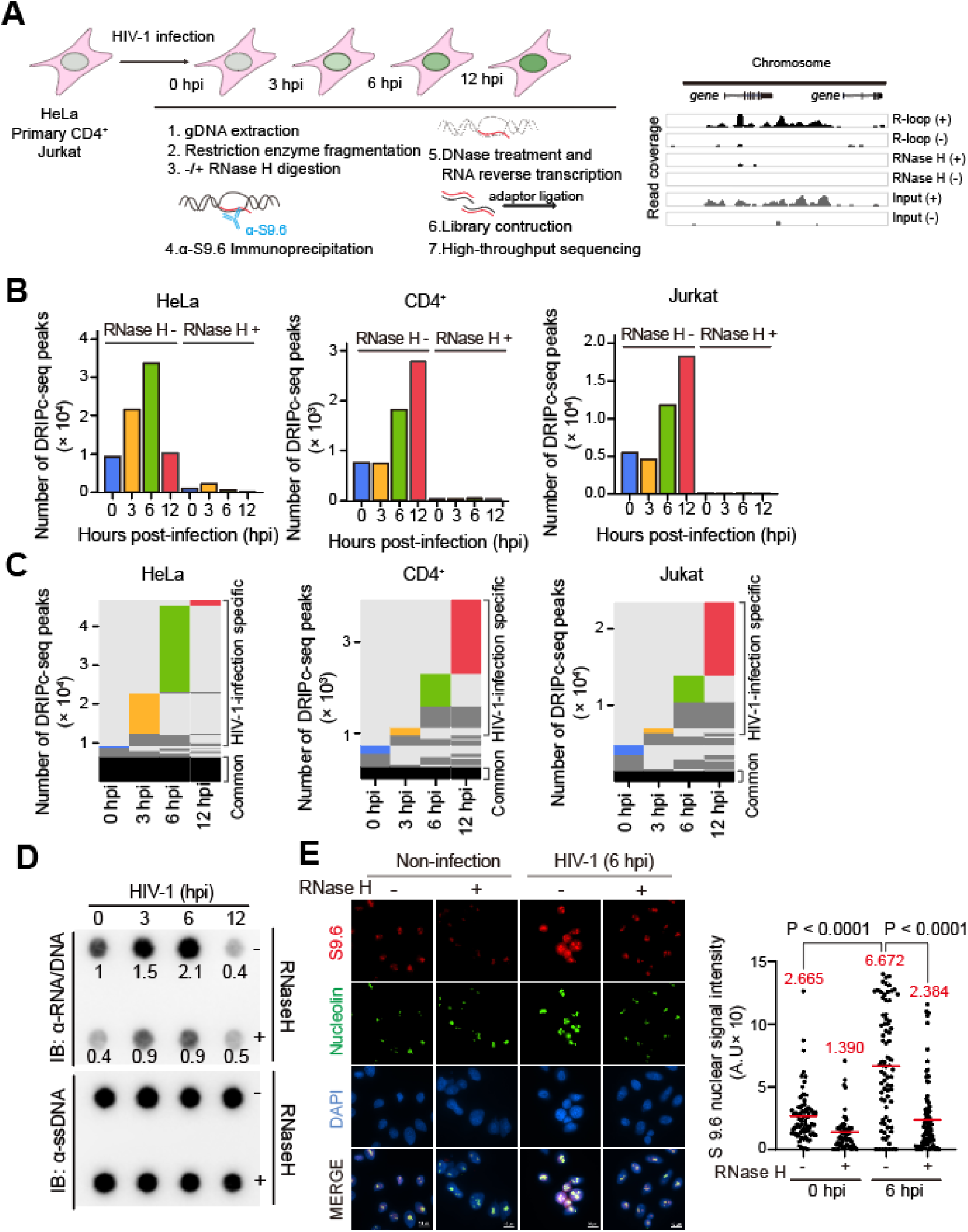
HIV-1 infection induces genomic R-loop accumulation in cells at early post-infection. (**A**) Summary of experimental design for DRIPc-seq in HeLa cells, primary CD4^+^ T cells and Jurkat cells infected with HIV-1. (**B**) Bar graphs indicating DRIPc-seq peak counts for HIV-1-infected HeLa cells, primary CD4^+^ T cells and Jurkat cells harvested at the indicated hours post infection (hpi). Pre-immunoprecipitated samples were untreated (−) or treated (+) with RNase H, as indicated. Each bar corresponds to pooled datasets from two biologically independent experiments. (**C**) All genomic loci overlapping a DRIPc-seq peak from HIV-1 infected HeLa cells, primary CD4^+^ T cells and Jurkat cells in at least one sample are stacked vertically; the position of each peak in a stack is constant horizontally across samples. Each hpi occupies a vertical bar, as indicated. Each bar corresponds to pooled datasets from two biologically independent experiments. Common peaks for all samples are represented in black, and in dark gray for those common for at least two samples. The lack of a DRIP signal over a given peak in any sample is shown in light gray. The sample-unique peaks are colored blue, yellow, green, and red at 0, 3, 6, and 12 hpi, respectively. (**D**) Dot blot analysis of the R-loop in gDNA extracts from HIV-1 infected HeLa cells with MOI of 0.6 harvested at the indicated hpi. gDNAs were probed with anti-S9.6. gDNA extracts were incubated with or without RNase H in vitro before membrane loading (anti-RNA/DNA signal). Fold-induction was normalized to the value of harvested cells at 0 hpi by quantifying the dot intensity of the blots and calculating the ratios of the S9.6 signal to the total amount of gDNA (anti-ssDNA signal). (**E**) Representative images of the immunofluorescence assay of S9.6 nuclear signals in HIV-1 infected HeLa cells with MOI of 0.6 harvested at 6 hpi. The cells were pre-extracted of cytoplasm and co-stained with anti-S9.6 (red), anti-nucleolin antibodies (green), and DAPI (blue). The cells were incubated with or without RNase H in vitro before staining with anti-S9.6 antibodies, as indicated. Quantification of S9.6 signal intensity per nucleus after nucleolar signal subtraction for the immunofluorescence assay. The mean value for each data point is indicated by the red line. Statistical significance was assessed using one-way ANOVA (n >53).

In addition to the DRIPc-seq data analysis, we used different biochemical approaches to examine R-loop accumulation after HIV-1 infection in HeLa cells. R-loop accumulation in HIV-1-infected cells was observed using DNA–RNA hybrid dot blots with the anti-S9.6 antibodies (Fig. 1D). Dot intensity significantly increased upon HIV-1 infection at 6 hpi and the enhanced R-loop signals on the dot blots of HIV-1-infected cells were highly sensitive to *in vitro* treatment with RNase H (Fig. 1D). This result is highly consistent with our DRIPc-seq data analysis of HIV-1-infected HeLa cells. Subsequently, we observed HIV-1-induced R-loops using an immunofluorescence assay by probing HIV-1-infected or non-infected control cells with S9.6 antibody at 6 hpi (Fig. 1E). After subtracting the nucleolar signal, the nuclear fluorescence signal associated with R-loops was significantly enhanced in cells infected with HIV-1 (Fig. 1E). We validated and quantified HIV-1-infection induced R-loop formation on the host genome in a genome-site-specific manner by using DRIP followed by real-time polymerase chain reaction (DRIP-qPCR). In this experiment, the S9.6 signal was determined for three and two HIV-1-induced-R-loop-positive (P1, P2, and P3) and –negative regions (N1 and N2), respectively, which were defined by DRIPc-seq data analysis (S2A-E Fig.). We detected significantly increased R-loop signals that were highly sensitive to RNase H treatment of pre-immunoprecipitates in the P1, P2, and P3 regions of HIV-1-infected cells at 6 hpi compared to those in the cells harvested at 0 hpi (S3A Fig.). However, the HIV-1-induced R-loop-negative regions N1 and N2, did not show significant R-loop accumulations (S3A Fig.).

Importantly, the R-loop signal was enriched even in cells infected with HIV-1 when the reverse transcription or integration of HIV-1 was blocked by enzyme inhibitors, such as nevirapine (NVP) or raltegravir (RAL) (S3B and S3C Fig.). This result indicates that the enrichment of R-loop signals in cells originates from the host genome, but not from DNA-RNA hybrid formation during the viral life cycle or transcriptional burst from integrated HIV-1 proviruses. In addition, we confirmed that nearly 100% of DRIPc-seq reads from HIV-1-infected HeLa, CD4^+^, and Jurkat T cells were aligned to the host cellular genome, but not to that of HIV-1 (S3D Fig.). Together, these data demonstrated that HIV-1 infection induce host genomic R-loop enrichment at early post-infection. This also suggests that HIV-1 infection regulates R-loop dynamics in host cells, possibly by inducing either new R-loop formation or stabilizing existing R-loops in the host cellular genome.

### R-loops accumulation after HIV-1 infection are widely distributed in both genic and non-genic regions

To investigate the distribution of cellular genomic R-loops during HIV-1 infection, we conducted a genome-wide analysis of the DRIPc-seq data. Unique DRIPc-seq peaks observed after HIV-1 infection were numerous and relatively long (Fig. 2A). This suggests that the R-loops induced by HIV-1 infection occupy a genomic region larger than that of the R-loops without HIV-1 infection. We observed a significant accumulation of R-loops over diverse genomic compartments at the hpi of HIV-1-infection induced R-loop formation (Fig. 2B). The presence of R-loops is often correlated with high transcriptional activity, and we found a significantly high proportion of DRIPc-seq peaks enrichment upon HIV-1 infection in the gene body regions (Fig. 2B). However, we also observed enrichment of HIV-1-infection induced DRIPc-seq peaks proportions mapped to intergenic or repeat regions, including short interspersed nuclear elements (SINEs), long interspersed nuclear elements (LINEs), and long terminal repeat (LTR) retrotransposons, where transcription is typically repressed (Fig. 2B). In addition, the proportion of DRIPc-seq peaks mapped to various genomic compartments remained consistent over the hours following HIV-1 infection (Fig. 2C). This suggests that HIV-1 infection does not induce R-loop enrichment at specific genomic features, but that R-loops accumulation after HIV-1 infection is widely distributed. Although the expression of repetitive elements is mostly repressed during normal cellular activities, HIV-1 infection could activate endogenous retroviral promoters (*30, 31*). To investigate whether R-loop induction in gene-silent regions is associated with transcriptome changes during HIV-1 infection, we performed RNA sequencing (RNA-seq) of HIV-1-infected HeLa cells at 0, 3, 6, and 12 hpi. Consistent with previous reports, we observed an increase in the expression levels of repetitive elements at later time points post-infection (S4A Fig. 12 hpi). In contrast, we found that the expression levels of SINEs, LINEs, and LTRs were even lower at both 3 and 6 hpi compared with those at 0 hpi whereas HIV-1-induced R-loops were significantly accumulated compared with those at 0 hpi (S4A Fig.). We further examined the expression profiles of genes containing the R-loop in HeLa cells. The expression profile of genes harboring HIV-1-induced R-loops in their gene bodies showed very weak correlations with the signals of DRIPc-seq peaks at 3 hpi (Pearson’s r = 0.21, P = 1.08 × 10^−84^; Fig. 2D) and at 6 hpi (Pearson’s r = –0.34, P = 2.40 × 10^−228^; Fig. 2D), which implies that the unique R-loop peaks upon HIV-1 infection do not engage in a transcriptional burst. In agreement with our DRIPc-seq and global RNA-seq data analysis, the expression level of the genes harboring HIV-1-infection induced R-loops, which were quantified by DRIP-qPCR (S3A Fig.), were not significantly affected by HIV-1 infection (S4B Fig. and Table S4). Together, our data demonstrate that host cellular R-loop accumulation upon HIV-1 infection is widely distributed in both genic and non-genic regions and is not necessarily correlate with the expression levels of the genes harboring the R-loops.

**Fig. 2.**
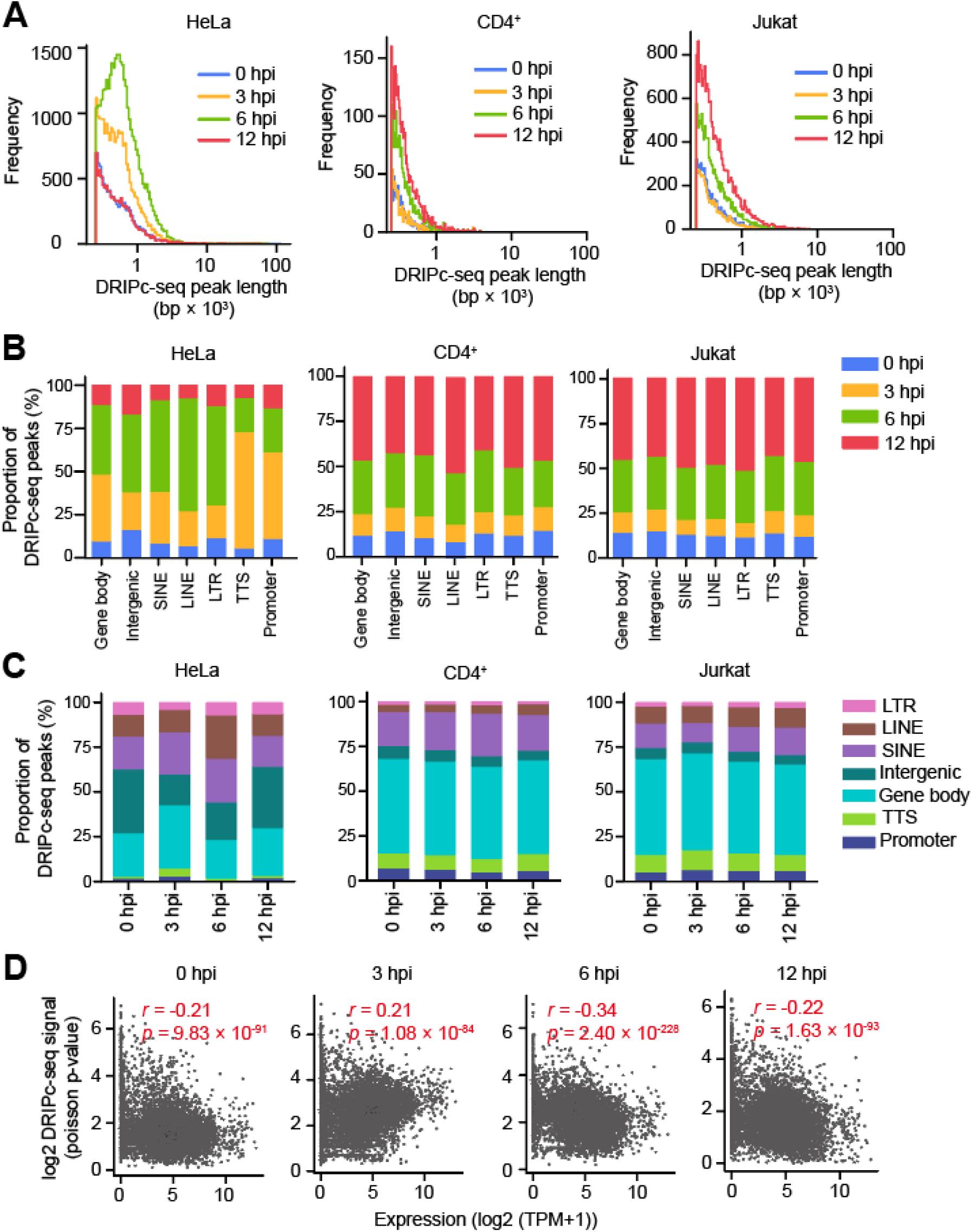
HIV-1-induced R-loops are enriched at both transcriptionally active and silent regions. (**A**) Distribution of DRIPc-seq peak lengths for HIV-1-infected HeLa cells, primary CD4^+^ T cells and Jurkat cells harvested at the indicated time points (blue, 0 hpi; yellow, 3 hpi; green, 6 hpi; red, 12 hpi). (**B**) Stacked bar graphs indicating the proportion of DRIPc-seq peaks mapped for HIV-1-infected HeLa cells, primary CD4^+^ T cells and Jurkat cells harvested at the indicated hpi over different genomic features. (**C**) Stacked bar graphs indicating the proportion of DRIPc-seq peaks mapped to indicated genomic compartments for HIV-1-infected HeLa cells, primary CD4^+^ T cells and Jurkat cells harvested at the 0, 3, 6, and 12 hpi. (**D**) Correlation between gene expression and DRIPc-seq signals of HIV-1-infected HeLa cells with MOI of 0.6 harvested at the indicated hpi. Statistical significance was assessed using Pearson’s r and p-values.

### HIV-1 integration sites are enriched at systemically induced sequence-specific R-loop regions in cell model

HIV-1 infection is completed by integrating the viral genome into the host’s through dynamic interactions with the host genome (*32*). In addition, as HIV-1 infection induces R-loop accumulation at early post infection hours, when the HIV-1 genome are imported into the nucleus and integration may initiate (*33–35*), we hypothesized that host genomic R-loops play a role in HIV-1 integration and possibly in integration site selection. To systemically and directly assess the relationship between host genomic R-loops and HIV-1 integration in a genome site-specific manner, we adapted and modified an elegantly designed episomal system that induces sequence-specific R-loops through DOX-inducible promoters (*17*). To most closely mimic the presence of the R-loop in the host cellular genome, we subcloned the R-loop-forming portion of the mouse gene encoding AIRN (mAIRN) (*18*) or the non-R-loop-forming ECFP sequence (*17*) with a DOX-inducible promoter into the piggyBac transposon vector and co-expressed piggyBac transposase in HeLa cells. The mAIRN gene sequence possesses a high GC skew that causes R-loop structure formation upon transcription by hybridizing newly synthesized RNA back to the template DNA strand, and the non-template DNA strand to remains looped out in a single-stranded form (*18, 36*). In contrast, the ECFP sequence possesses a low GC skew that does not form a stable R-loop at the site of transcription. Thus, it was used as an R-loop-forming negative sequence. These R-loop forming (mAIRN) or non-R-loop forming (ECFP) sequences are nonhuman sequences. Therefore, our cell model allowed us to induce and quantify R-loop formation at designated genomic regions and distinguish R-loop formation from endogenous R-loops on the cellular genome, which are not sequence-specific and impossible to control for induction. Moreover, using this system we can quantify R-loop-associated site-specific HIV-1 integration events at designated regions, which can also be distinguished from HIV-1 integration events at endogenous host genomic loci. We designated the pool of cells with the R-loop forming sequence (mAIRN) inserted into its genome as “pgR-rich (piggyBac R-loop rich)” cell line and the pool of cells with the non-R-loop forming sequence (ECFP) inserted into its genome as “pgR-poor (piggyBac R-loop poor)” cell line (Fig. 3A).

**Fig. 3.**
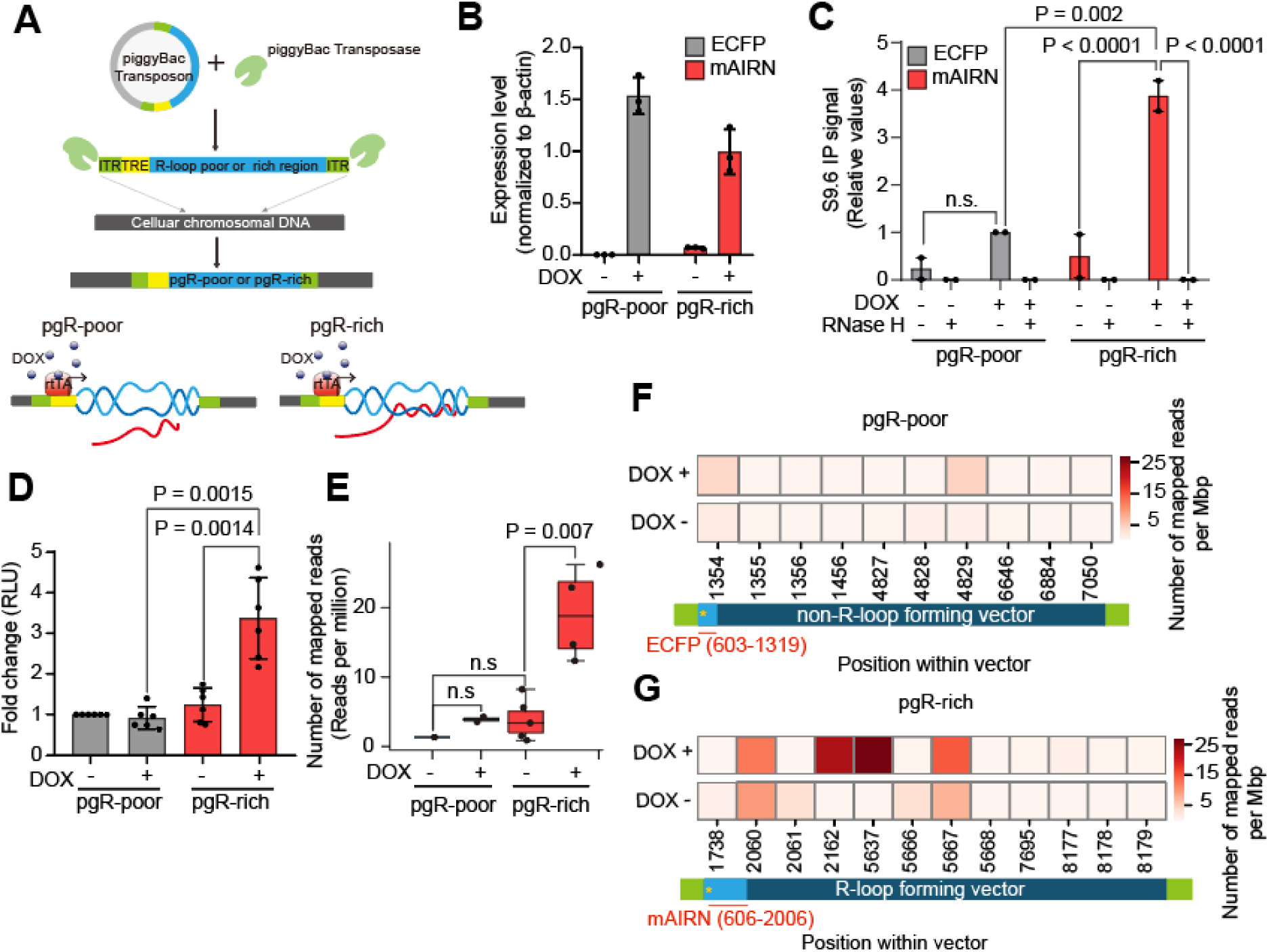
R-loop inducible cell line model directly addresses R-loop-mediated HIV-1 integration site selection. (**A**) Summary of the experimental design for R-loop inducible cell lines, pgR-poor and pgR-rich. (**B**) Gene expression of ECFP (gray) and mAIRN (red), as measured using RT-qPCR in pgR-poor or pgR-rich cells. Where indicated, the cells were incubated with 1 µg/ml DOX for 24 h. Gene expression was normalized relative to *β-actin*. Data are presented as the mean ± SEM, n = 3. (**C**) DRIP-qPCR using the anti-S9.6 antibody against ECFP and mAIRN in pgR-poor or pgR-rich cells. Where indicated, the cells were incubated with 1 µg/ml DOX for 24 h. Pre-immunoprecipitated samples were untreated or treated with RNase H as indicated. Values are relative to those of DOX-treated (+) RNase H-untreated (−) pgR-poor cells. Data are presented as the mean ± SEM; statistical significance was assessed using two-way ANOVA (n = 2). (**D**) Bar graphs indicate luciferase activity at 48 hpi in pgR-poor or prR-rich cells infected with 100ng/p24 capsid antigen of luciferase reporter HIV-1 virus per 1× 10^5^ cells/mL. Data are presented as the mean ± SEM; P values were calculated using one-way ANOVA (n = 6). (**E**) Box graph indicating the quantified HIV-1 integration site sequencing read count across pgR-poor and pgR-rich transposon sequences in untreated (−) or DOX-treated (+) pgR-poor or pgR-rich cell line infected with 100ng/p24 capsid antigen of luciferase reporter HIV-1 virus per 1× 10^5^ cells/mL. Each bar corresponds to pooled datasets from three biologically independent experiments (n =3). In each boxplot, the centerline denotes the median, the upper and lower box limits denote the upper and lower quartiles, and the whiskers denote the 1.5 × interquartile range. Statistical significance was assessed using a two-sided Mann–Whitney U test. (**F** and **G**) Heat maps representing the number of HIV-1 integration-seq mapped read across pgR-poor (**F**) or pgR-rich (**G**) transposon sequence in untreated (-) or DOX-treated (+) pgR-poor (**F**) or pgR-rich (**G**) cell line. Each rectangular box corresponds to the pooled the number of HIV-1 integration-seq mapped read from three biologically independent experiments (n =3) at the indicated position within pgR-poor (**F**) or pgR-rich (**G**) transposon vector. Each light blue box represents actual position of R-loop forming or non-R-loop forming sequence (ECFP or mAIRN) and the yellows stars indicate TRE promoter position within vector.

A similar number of the copies of the piggyBac transposon were successfully delivered to the genome of each cell line (S5A Fig.) and DOX treatment strongly induced the transcriptional activity of mAIRN or ECFP without affecting the transcription of endogenous loci in either cell line (S5B and S5C Fig.). Although the transcription of mAIRN or ECFP was strongly induced by DOX treatment, the activity did not exceed that of the endogenous loci in either cell line (S5D and S5E Fig.). Although the two cell lines showed comparable levels of DOX-inducible transcriptional activity at the designated sequences (Fig. 3B), only the pgR-rich cells exhibited robust RNase H-sensitive stable R-loop formation upon DOX treatment (Fig. 3C, mAIRN). In contrast, R-loops were weakly formed in the pgR-poor cells where a non-R-loop-forming sequence (ECFP) was inserted into the genome (Fig. 3C, ECFP).

To examine whether the formation of ‘extra’ R-loops in the host genome influence HIV-1-infection in host cells, we infected both cell lines with VSV-G-pseudotyped HIV-1-luciferase viruses and examined the luciferase activity. Notably, we found that pgR-rich cells showed significantly higher luciferase activity only when R-loops were induced by DOX treatment, whereas pgR-poor cells showed comparable luciferase activity regardless of transcriptional activation by DOX treatment (Fig. 3D). We sequenced HIV-1 integration sites in HIV-1-infected pgR-poor and pgR-rich cells to directly quantify site-specific integration events in sequence-specific R-loop regions. Remarkably, integration events were significantly higher in pgR-rich cells only when R-loops were induced by DOX treatment (Fig. 3E). However, the number of HIV-1 integration-seq mapped reads within the non-R-loop forming sequence in pgR-poor cells remained very low, even after transcriptional activation by DOX treatment (Fig. 3E). HIV-1 integration was enriched in the vicinity of R-loop forming regions in the pgR-rich cell line upon DOX treatment, but the enrichment was not observed in pgR-poor cells that did not form stable R-loops, even after transcriptional activation by DOX treatment (Figs. 3F and 3G). This cell-based R-loop-inducing system with independent control over transcription and R-loop formation enabled the direct measurement of HIV-1 integration events at defined R-loop regions. The results indicated that host genomic R-loops are preferred by HIV-1 integration. Moreover, our data suggest that transcriptional activity itself is not sufficient for HIV-1 integration site determination, but that the formation of R-loops accounts for HIV-1 integration site selection.

### Host genomic R-loop regions are frequently targeted by HIV-1 integration

We attempted to further validate the relationship between R-loops and HIV-1 integration site selection by global analysis of HIV-1 integration sites in the endogenous genomic regions of HIV-1 infected host cells. We performed HIV-1 integration site sequencing in HIV-1 infected HeLa cells, CD4^+^, and Jurkat T cells, and analyzed the sequencing data combined with our DRIPc-seq data. We counted and compared the number of successfully integrated proviruses in the R-loop regions (the combined genomic regions within 30-kb windows centered on DRIPc-seq peaks from 0, 3, 6, and 12 hpi) to those in non-R-loop-forming regions (the total genomic regions outside of the 30-kb windows centered on DRIPc-seq peaks). Notably, we detected approximately three to four times more integration in the R-loop regions than in other genomic regions without R-loops in HeLa cells, CD4^+^, and Jurkat T cells (Fig. 4A). Notably, HIV-1 integration sites preferred the centeral and nearby areas of the R-loops regions (Fig. 4B). We observed biases in HIV-1 integration in HIV-1-induced R-loop-positive regions, P1-P3, where showed highly induced R-loop signals upon HIV-1 infection in DRIPc-seq analysis and DRIP-qPCR (Fig. 4C). In contrast, HIV-1 integration sites were not detected in the R-loop-negative regions N1 and N2 (Fig. 4D). Overall, our results from bioinformatics analysis using different types of naïve host cells infected with HIV-1 are consistent with the idea that the virus has a preference for targeting R-loop-forming regions for integration (Fig. 3), suggesting that R-loops are an important component of the host genomic environment for HIV-1 integration site determination.

**Fig. 4.**
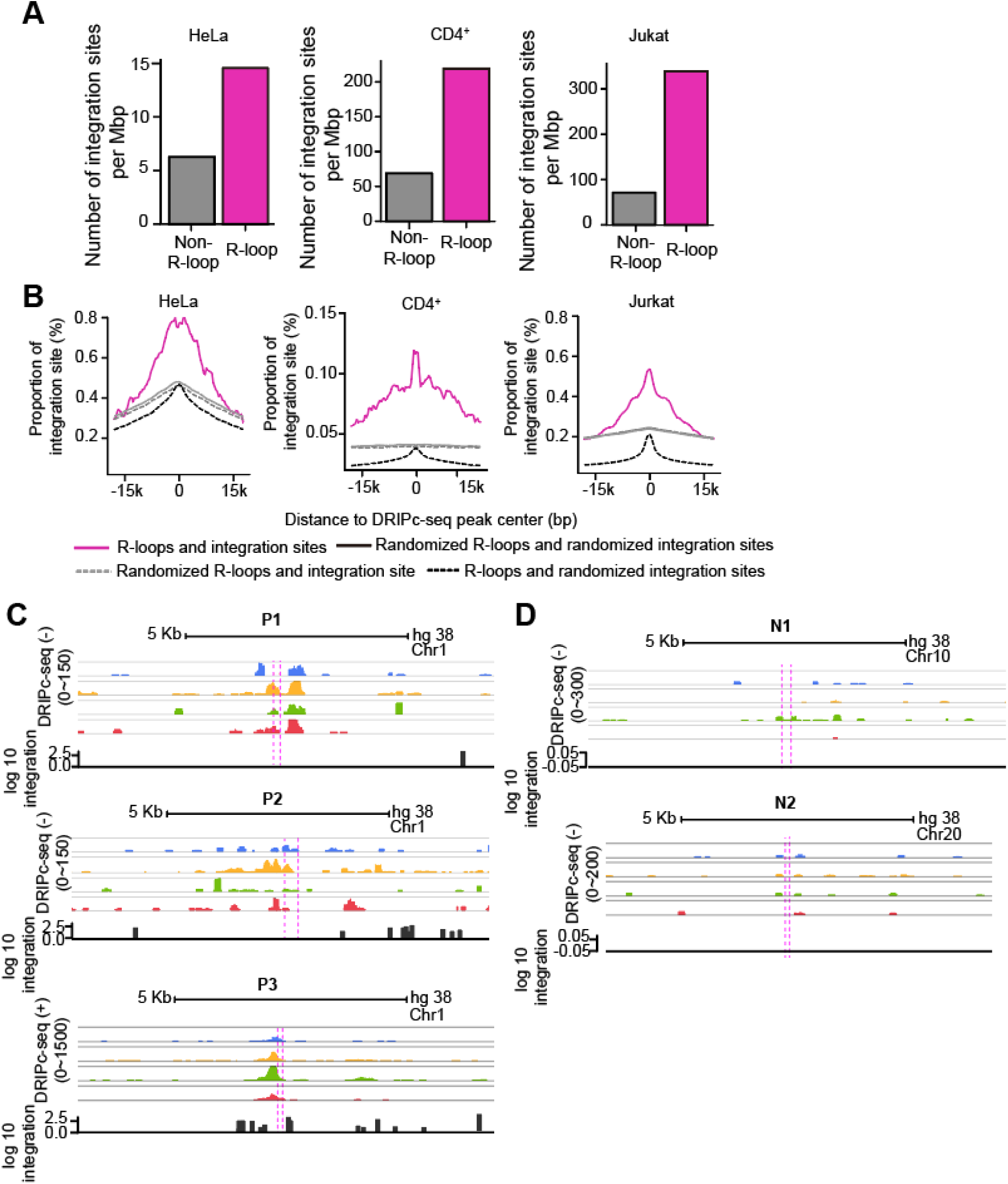
HIV-1 prefers host genomic R-loop regions for its viral cDNA integration. (**A**) Bar graphs showing quantified number of HIV-1 integration sites per Mb pairs in total regions of 30-kb windows centered on DRIPc-seq peaks from HIV-1 infected HeLa cells, primary CD4^+^ T cells and Jurkat cells (magenta) or non-R-loop region in the cellular genome (gray). (**B**) Proportion of integration sites within the 30-kb windows centered on DRIPc-seq peaks (magenta solid lines) or randomized DRIPc-seq peaks (gray dotted lines). Control comparisons between randomized integration sites with DRIPc-seq peaks and randomized DRIPc-seq peaks are indicated by black dotted lines and gray solid lines, respectively. (**C** and **D**) Superimpositions of HIV-1-induced R-loop positive chromatin regions, P1-P3 (**C**), and HIV-1-induced R-loop negative chromatin regions, N1 and N2 (**D**), on DRIPc-seq (blue, 0 hpi; yellow, 3 hpi; green, 6 hpi; red, 12 hpi) and number of mapped read of HIV-1 integration-seq (integration, black). Magenta dotted lines represent primer binding sites in qPCR following DRIP.

### HIV-1 integrase proteins physically interacts with R-loops on the host genome

The HIV-1 intasome is tethered to the host genome for viral cDNA integration. Intasomes consist of HIV-1 viral cDNA and the HIV-1 integrase proteins. We observed that HIV-1 was preferentially integrated into the R-loops regions in the host genome; thus, we hypothesized that the HIV-1 integrase protein could directly bind and be recruited to the genomic R-loops. To test this hypothesis, we investigated whether HIV-1 integrase proteins have a physical binding affinity for nucleic acid substrates with an R-loop structure. Although HIV-1 integrases are DNA and RNA binding proteins (*37, 38*), their ability to bind to a three-stranded nucleic acid structure that is composed of a DNA-RNA hybrid like R-loop has not been investigated. We carried out an *in vitro* protein-nucleic acid binding assay by electrophoretic mobility shift assay (EMSA) with Sso7d-tagged HIV-1 integrase (E152Q) recombinant proteins and diverse structures of nucleic acid substrates including R-loops and simple dsDNA duplexes. In this experiment, we used HIV-1 integrase protein with an active site amino acid substitution, E152Q, to prevent any undesirable alternation in nucleic acid substrates by enzymatic activities of integrase proteins, such as 3’-processing. Notably, nucleic acid substrate consisted of an R-loop structure bound to HIV-1 integrase proteins with higher binding affinity than that of the simple dsDNA duplex (Fig. 5A). Additionally, the R-loop composing forms of nucleic acid structures, such as RNA-DNA hybrid with exposed ssDNA (R:D+ssDNA) and RNA-DNA hybrid (hybrid), also showed high binding affinity to integrases (S6A Fig. and Fig. 5A).

**Fig. 5.**
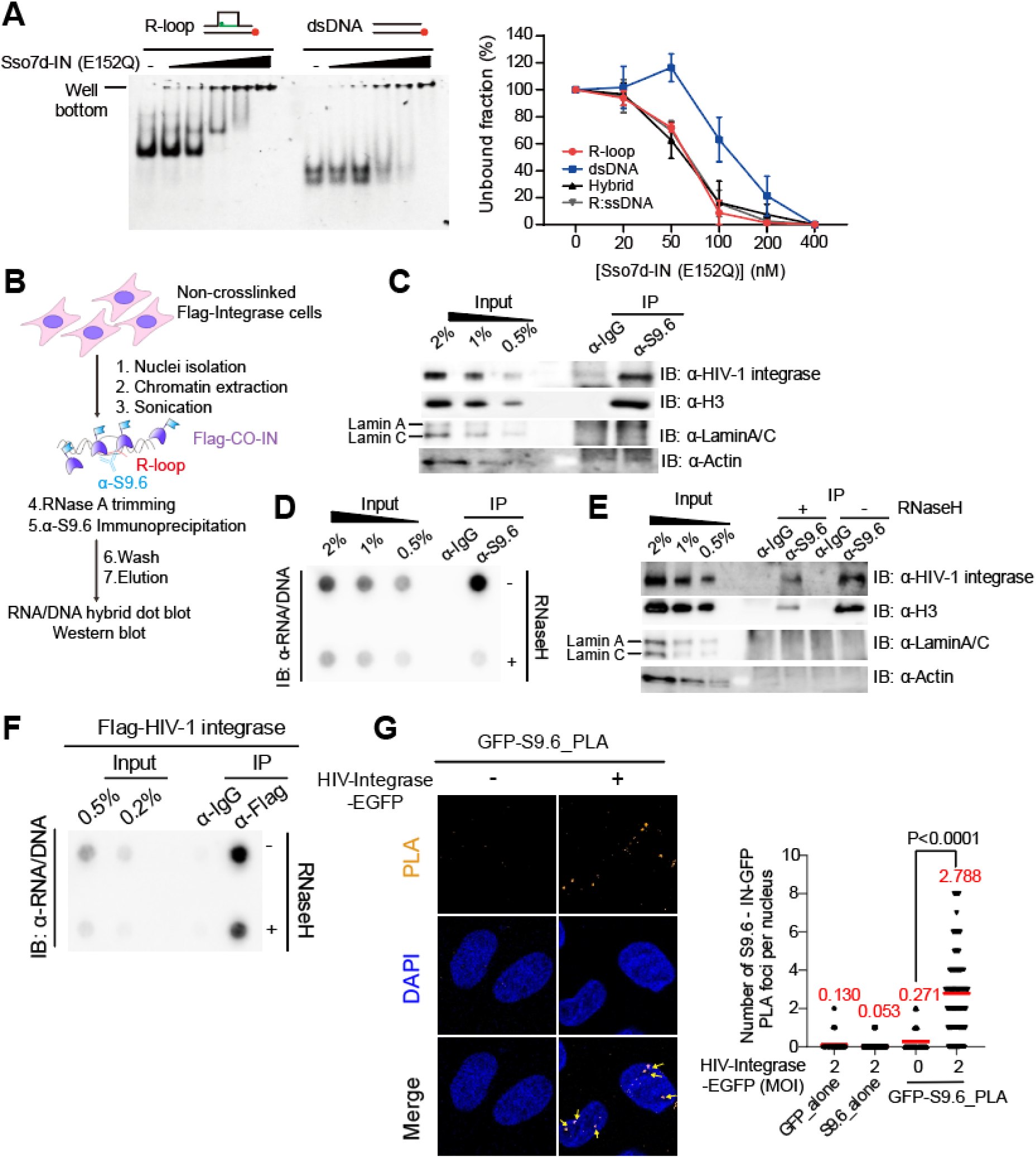
HIV-1 integrase proteins directly bind to host genomic R-loops. (**A**) Representative gel images for EMSA of Sso7d-tagged HIV-1-integrase (E152Q) with R-loop and dsDNA, 10 nM nucleic acid substrate was incubated with Sso7d-tagged HIV-1-integrase (E152Q) at 0 nM, 20 nM, 50 nM, 100 nM, 200 nM, and 400 nM (left). Unbound fraction were quantified for EMSA of Sso7d-tagged HIV-1-integrase (E152Q) with different types substrates (R-loop, dsDNA, R-loop, R:D+ssDNA and Hybrid). Data are presented as the mean ± SEM, n = 3 (right). (**B**) Summary of the experimental design for R-loop immunoprecipitation using S9.6 antibody in FLAG-tagged HIV-1 integrase protein-expressing HeLa cells. (**C**) Western blotting for HIV-1 integrase protein, H3, and LaminA/C of DNA–RNA hybrid immunoprecipitation using the S9.6 antibody. (**D**) **and** (**E**) HeLa gDNA input was either untreated (−) or treated (+) with RNase H before enrichment for DNA–RNA hybrids using the S9.6 antibody. gDNA–RNA hybrids were incubated with nuclear extracts depleted of DNA–RNA hybrids with RNase A followed by S9.6 immunoprecipitation. DNA–RNA hybrid dot blot (**D**) and western blot of DNA–RNA hybrid immunoprecipitation, probed with the indicated antibodies (**E**). (**F**) DNA–RNA hybrid dot blot of FLAG antibody-immunoprecipitated nucleic acid extracts. Where indicated, nucleic acid extracts were untreated (−) or treated (+) with RNase H before probing with the S9.6 antibodies. (**G**) Representative images of the proximity-ligation assay (PLA) between GFP and S9.6 antibodies in HIV-IN-EGFP virion-infected HeLa cells at 6 hpi. Cells were subjected to PLA (orange) and co-stained with DAPI (blue). PLA puncta in the nucleus are indicated by the yellow arrows. Quantification analysis of number of PLA foci per nucleus (left). GFP_alone and S9.6_alone were used as single-antibody controls from HIV-IN-EGFP virion-infected HeLa cells (right). The mean value for each data point is indicated by the red line. P value was calculated using a two-tailed unpaired *t*-test (n > 50).

We validated the interaction between cellular genomic R-loops and HIV-1 integrase proteins by DNA–RNA hybrid immunoprecipitation using S9.6 antibodies against FLAG-tagged HIV-1 integrase-expressing HeLa cells (Fig. 5B). Under our experimental conditions, R-loops were reproducibly immunoprecipitated (S6B Fig.) and HIV-1 integrase proteins co-immunoprecipitated with the R-loops (Fig. 5C). DNA–RNA hybrids also co-immunoprecipitated with the positive control H3 (*39*) but not with the negative controls LaminA/C and Actin (*39*) (Fig. 5C). To verify the specificity of our co-immunoprecipitation results for R-loops and HIV-1 integrases, we performed DNA–RNA hybrid immunoprecipitation with RNase H treatment (S6C Fig.). The S9.6 signal of the immunoprecipitated nucleic acids was highly sensitive to RNase H treatment of the pre-immunoprecipitates (Fig. 5D). Accordingly, the blotting signal of the co-immunoprecipitated HIV-1 integrase and H3 proteins was significantly reduced upon RNase H treatment (Fig. 5E). We performed reciprocal immunoprecipitation using an anti-FLAG monoclonal antibody and detected the immunoprecipitated R-loops using dot blot analysis with anti-S9.6. R-loops, which were immunoprecipitated using HIV-1 integrase, and the S9.6 signal of immunoprecipitated nucleic acids was highly sensitive to RNase H treatment (Fig. 5F and S6D Fig.). Subsequently, we examined the interaction between the R-loops and HIV-1 integrase using a proximity-ligation assay (PLA) in HIV-1-infected cells. We used two antibodies: one that binds to the R-loops (anti-S9.6) and another that binds to GFP-tagged HIV-1 integrase. We detected PLA signals in cells infected with HIV-IN-EGFP virions (*34*) and non-infected control cells. PLA signals in non-infected cells were comparable to those in S9.6-alone and GFP-alone single antibody-negative controls; however, PLA signals significantly increased upon HIV-1 infection (Fig. 5G and S6E Fig.). Our data suggest that the HIV-1 frequently targets R-loop-rich regions for viral genome integration by physically binding of HIV-1 integrase proteins to R-loop structures on the host genome.

## Discussion

In this study, we found that HIV-1 preferentially integrated into regions rich in R-loops, suggesting that R-loops are novel host factors that contribute to HIV-1 integration site selection. In our bioinformatics analysis, host cellular R-loops were induced by VSV-G-pseudotyped HIV-1 infection and widespread in host genomic regions. Using our R-loop-inducible cell models, R-loop formation, and not necessarily the transcriptional activity itself, was found to be important for HIV-1 integration site determination. In addition, HIV-1 integrase proteins favor physical binding with R-loops *in vitro*, and interact with host genomic R-loops in HIV-1-infected cells. These results demonstrate that HIV-1 exploits and frequently targets the host genomic R-loop regions for successful integration and infection.

One possible explanation for why HIV-1 integration shows a preference for host genomic R-loop regions is that the R-loop structure may drive dynamics in the genomic environment and the spatial organization of the genome, resulting in increased accessibility of HIV-1 intasome binding to the target host genomic region. The R-loops display enhancer and insulator chromatin states, that can act as distal regulatory elements by recruiting diverse chromatin binding factors (*22*). This not only allows R-loops to drive dynamics in the genome, but also possibly drives R-loop-mediated integration over long-range genomic regions. R-loop regions exhibit increased chromatin accessibility. In the cellular genome, these structures relieve superhelical stresses and are often associated with open chromatin marks and active enhancers (*22, 23*), which are also distributed over HIV-1 integration sites (*6*). In the case of transcription-induced R-loop formation, a guanine-quadruplex (G4) structure can be generated in the non-template DNA strand of the R-loop, which is another contributor to genome architecture (*24*). A recent study showed that G4 DNA can influence both productive and latent HIV-1 integration, as well as the potential for reactivation of latent proviruses (*25*).

Another possible explanation for the preference of HIV-1 integration for host genomic R-loops is that R-loops may play a collaborative role with host factors governing the HIV-1 integration site selection. Cellular R-loops are recognized and regulated by numerous cellular proteins (*39, 40*). LEDGF/p75 (*9, 13, 41*) and CPSF6 (*7, 9*) are two decisive host factors that direct HIV-1 integration by interacting with integrase or trafficking the viral preintegration complex towards the nuclear interior (*7, 9*). In fact, these host factors have recently been identified as potential R-loop-binding proteins in DNA–RNA interactome analysis (*39*) and R-loop proximity proteomics (*40*), respectively. R-loops are tightly regulated by DNA damage response proteins (*42*) and the DNA repair machinery plays an important role in HIV-1 integration process (*32*). For example, the Fanconi anemia pathway (*43, 44*), a well-known R-loop regulatory pathway, has been recently proposed as an HIV-1 integration regulatory factor exploited by HIV-1 (*45*). Considering theses previous studies and our current findings, we propose R-loops as another putative host factor driving HIV-1 integration site determination, and a possible intermediate regulator of HIV-1 integration site selection by such host proteins.

Our data showed that HIV-1 integrase proteins physically interacted with genomic R-loops *in vitro* and in cells. Recent advancements in cryogenic electron microscopy (cryo-EM) technology have revealed the conformational characteristics of the target DNA during retroviral integration (*27, 28*). During retroviral integration, the target DNA undergoes a transition of its conformation from B-form to A-form. R-loops, which represent intermediates between the B-form DNA and A-form RNA conformations (*27*), may have an intrinsic preferential binding ability to retroviral intasomes over other nucleic acid structures.

Viruses often take advantage of various host factors, and targeting the viral components that manipulate the host cellular environment can be an effective strategy for antiviral therapy. Our study showed the host genomic R-loops significantly accumulate shortly after HIV-1 infection. Thus, it is possible that virion-associated HIV-1 proteins are responsible for inducing these R-loops. For instance, the HIV-1 accessory protein Vpr causes genomic damage (*46*) and transcriptomic changes during the early stages post infection(*47*), both of which can lead to *in cis* and *in trans* R-loop formation (*16*). Another HIV-1 accessory protein, Vif, counteracts the host antiviral factor, APOBEC3 (*48, 49*), which was recently found to regulates cellular R-loop levels (*50*). Identifying the HIV-1 components responsible for inducing host cellular R-loops, elucidating the mechanism by which they induce genome-wide R-loop formation, and contributing to successful viral integration into selective genomic regions, are areas for further research.

Although most HIV-1 integration occurs in genic regions (*4, 6*), HIV-1 proviruses are also found in non-genic regions (*51*) and understanding these “transcriptionally silent” proviruses is critical for developing strategies to completely eliminate HIV-1. In HIV-1 elite controllers, who suppress viral gene expression to undetectable levels, HIV-1 proviruses in heterochromatic regions are not eliminated but selected by the immune system (*5*). Moreover, proviruses with lower expression level can persist in the host genome even during antiretroviral therapy (*4*). However, the mechanism by which HIV-1 targets gene-silent regions for “invisible” integration remains unclear. Our study has revealed that R-loops are enriched in both genic and non-genic regions during HIV-1 infection, and that the virus preferentially targets these R-loops for integration. We propose that R-loops, particularly those enriched in non-genic regions, may represent the mechanism by which the virus achieves “invisible” and permanent infection.

## Materials and methods

### Cell culture

HeLa and HEK293T cells were cultured in Dulbecco’s modified Eagle’s medium (Gibco) supplemented with 10% (v/v) fetal bovine serum (FBS, Cytiva), antibiotic mixture (100 units/ml penicillin–streptomycin, Gibco), and 1% (v/v) GlutaMAX-I (Gibco). Jurkat cells were cultured in Roswell Park Memorial Institute (RPMI) 1640 medium (ATCC) supplemented with 10% (v/v) FBS (Cytiva). Cells were incubated at 37°C and 5% CO_2_.

### Virus production and infection

VSV-G-pseudotyped HIV-1 virus stocks were prepared by performing standard polyethylenimine-mediated transfection of HEK293T monolayers with pNL4-3 ΔEnv EGFP (NIH AIDS Reagent Program 11100) or pNL4-3. Luc.R-E (NIH AIDS Reagent Program, 3418) along with pVSV-G at a ratio of 5:1. HIV-IN-EGFP virions were produced as previously described (*34*) by performing polyethylenimine-mediated transfection of HEK293T cells with 6 µg of pVpr-IN-EGFP, 6 µg of HIV-1 NL4-3 non-infectious molecular clone (pD64E; NIH AIDS Reagent Program 10180), and 1 µg of pVSV-G. The cells were incubated for 4 h before the medium was replaced with fresh complete medium. Virion-containing supernatants were collected after 48 h, filtered through a 0.45-µm syringe filter, and pelleted using the Lenti-X Concentrator (631232; Clontech) according to the manufacturer’s instructions. The multiplicity of infection (MOI) of virus stocks was determined by transducing a known number of HeLa cells with a known amount of virus particles and then counting GFP-positive cells using flow cytometry. For luciferase reporter HIV-1 virus, the HIV-1 p24 antigen content in viral stock were quantified using the HIV1 p24 ELISA kit (Abcam, ab218268), according to the manufacturer’s instruction. For virus infection, HeLa cells were seeded at a density of 0.5–4 × 10^5^ cells/mL on the day before infection. The culture medium was replaced with fresh complete culture medium 2 hpi. The infected cells were washed twice with PBS and harvested at the indicated time points. Jurkat cells were seeded at a density of 1× 10^6^ cells/mL and inoculated with 300ng/p24 capsid antigen. The plates were centrifuged at 1000 *g* at 30°C for 1 h. The medium was replaced with fresh RPMI 2 h after infection.

### Primary cell isolation, culture, T cell activation, and infection

For CD4^+^ T cells isolation, human PBMC (ST70025, STEMCELL Technologies) was mixed and incubated with MACS CD4 MicroBeads (130-045-101, Miltenyi Biotec) and FITC-conjugated mouse anti-CD4 (561005, BD Bioscience) according to the manufacturer’s instructions. Then the CD4^+^ T cells were enriched by using LS Columns (130-042-401, Miltenyi Biotec) and MidiMACS Separator (130-042-302, Miltenyi Biotec). The efficiency of magnetic separation was analyzed by using Flow-Activated Cell Sorter Canto II (BD Bioscience) and Flowjo software (Flowjo).

CD4^+^ T cells were cultured in Roswell Park Memorial Institute (RPMI) 1640 medium (Gibco), supplemented with 10% (v/v) fetal bovine serum (FBS, Cytiva), antibiotic mixture (100 units/ml penicillin–streptomycin, Gibco), 1% (v/v) GlutaMAX-I (Gibco), and 20 ng/ml of IL-2 (PHC0026, Gibco), left in resting state or activated with Dynabeads Human T-Activator CD3/CD28 (1161D, Thermo Fisher Scientific) for 72 h. CD4^+^ T cells activation efficiency was assessed by staining cells with FITC-conjugated mouse anti-CD25 (340694, BD Bioscience) and APC-conjugated mouse anti-CD69 (130-114-046, Miltenyi Biotec) and using Flow-Activated Cell Sorter Canto II (BD Bioscience) and Flowjo software (Flowjo).

Purified and activated CD4^+^ T cells were seeded at a density of 1× 10^6^ cells/mL and inoculated with 600ng/p24 capsid antigen in presence of polybrene. The plates were centrifuged at 1000 *g* at 30°C for 1 h. The medium was replaced with fresh RPMI 2 h after infection.

### DRIP-qPCR

DRIP was performed as described for the construction of the DRIPc-seq library. After the elution of isolated complexes, nucleic acids were purified using the standard phenol-chloroform extract method and used for qPCR. S6 Table presents details of the primer sequences used for DRIP-qPCR analysis.

### RNA-seq library construction

For RNA-seq, HeLa cells were infected with VSV-G-pseudotyped HIV-1 NL4-3 ΔEnv EGFP virus at a MOI of 0.6 and harvested at 0, 3, 6, and 12 hpi. Sequencing was performed with biological replicates. Total RNA was extracted using TRIzol reagent (Invitrogen), according to the manufacturer’s instructions. An mRNA sequencing library was constructed using Illumina adaptors harboring p5 and p7 sequences and Rd1 SP and Rd2 SP sequences. Sequencing was performed using the HiSeq2500 system (Illumina).

### Luciferase assay

HeLa cells infected with VSV-G-pseudotyped pNL4-3.Luc.R-E HIV-1 viruses were harvested at 48 hpi, and luminescence was measured using the Dual-Luciferase Reporter Assay System (Promega) according to the manufacturer’s instructions. Briefly, 250 μl of passive lysis buffer was used to lyse cells for each sample, 20 μl of the lysate was mixed with 100 μl of the Luciferase Assay Reagent II, and the luminescence of firefly luciferase was measured using a microplate luminometer (Berthold). The luminescence signal were normalized with total protein content, measured by BCA assay.

### Quantitative real-time PCR (qPCR)

For RT (reverse transcription)-qPCR, 1 μg of RNA was reverse-transcribed using the ReverTra Ace qPCR RT Kit (TOYOBO) following the manufacturer’s instructions. For qPCR, DNA extracts were prepared using a DNA purification kit (Qiagen, 51106) according to the manufacturer’s instructions. Equivalent amounts of purified gDNA from each sample were analyzed using qPCR. qPCR was performed using TOPreal qPCR PreMIX (Enzynomics, RT500M). The reactions were performed in duplicate or triplicate for technical replicates. PCR was performed using the iCycler iQ real-time PCR detection system (Bio-Rad). All the primers used for qPCR are listed in S6 Table.

### DRIPc-seq library construction

DRIP followed by library preparation, next-generation sequencing, and peak calling were performed as described earlier (*29*). Briefly, the corresponding cells were harvested and their gDNA was extracted. The extracted nucleic acids were fragmented using a restriction enzyme cocktail with BsrB I (NEB, R0102S), HindIII (NEB, R0136L), Xba I (NEB, R0145L), and EcoRI (NEB, R3101L) overnight at 37°C. Half of the fragmented nucleic acids were digested with RNase H (New England Biolabs) overnight at 37°C to serve as a negative control. The digested nucleic acids were cleaned using standard phenol-chloroform extraction and resuspended in DNase/RNase-free water. DNA–RNA hybrids were immunoprecipitated from total nucleic acids using mouse anti-DNA–RNA hybrid S9.6 (Kerafast, ENH001) DRIP binding buffer and incubated overnight at 4°C. Dynabeads Protein A (Invitrogen, 10001D) was used to pull down the DNA-antibody complexes by incubation for 4 h at 4°C. The isolated complexes were washed twice with DRIP binding buffer before elution. For elution, the isolated complexes were incubated in an elution buffer containing proteinase K for 45 min at 55 °C. Subsequently, DNA was purified using the standard phenol-chloroform extract method and subjected to DNase I (Takara, 2270 B) treatment and reverse transcription for DRIPc-seq library construction. DRIPc-seq was performed in biological replicates. S5 Table shows details of the oligonucleotides used for DRIPc-seq library construction. DRIPc-seq libraries were analyzed using 150 bp paired-end sequencing on a HiSeqX Illumina instrument.

### Immunofluorescence microscopy

For immunofluorescence assays of S9.6 nuclear signals, when indicated, the cells were pre-extracted with cold 0.5% NP-40 for 3 min on ice. Cells were fixed with 100% ice-cold methanol for 10 min on ice and then incubated with 100% ice-cold acetone for 1 min. The slides were washed three times with 1× PBS and incubated with or without 60 U/mL RNase H (M0297S, NEB) at 37°C for 36 h or left untreated. The slides were subsequently briefly rinsed thrice with 2% BSA/0.05% Tween (in PBS) and incubated with mouse anti-DNA–RNA hybrid S9.6 (Kerafast, ENH001; 1:100) and rabbit anti-nucleolin (Abcam, ab22758; 1:300) in 2% BSA/0.05% Tween (in PBS) for 4 h at 4°C. The slides were then washed three times with 2% BSA/0.05% Tween (in PBS) and incubated with goat anti-rabbit AlexaFluor-488-conjugated (Invitrogen, A-11008) and goat anti-mouse AlexaFluor-568-conjugated (Molecular Probes, A11004) secondary antibodies (1:200) for 2 h at room temperature. The slides were then washed three times with 2% BSA/0.05% Tween (in PBS) and mounted using the ProLong Gold AntiFade reagent (Invitrogen). Images were obtained using an inverted microscope Nikon Eclipse Ti2, equipped with a 1.45 numerical aperture, plan apochromat lambda 100× oil objective, and an scientific complementary metal–oxide–semiconductor camera (Photometrics prime 95 B 25 mm). For each field of view, images were obtained with DAPI395, GFP488, and Alexa594 channels using the NIS-Elements software. For quantification analysis, binary masks of nuclei and nucleoli were generated using the ROI manager and auto local thresholding using the ImageJ software. The intensity of nuclear signals for DNA–RNA hybrids and nucleolin was then quantified. The final DNA–RNA hybrid signals in the nucleus were calculated by subtracting the nucleolin signals from the DNA–RNA hybrid signals.

### pgR-rich and –poor cell line generation with piggyBac transposition

We adapted and modified an elegantly designed episomal system that induces defined R-loops with controlled transcription levels (*17*) for R-loop-forming or non-R-loop-forming sequence subcloning into the piggyBac transposon vector. HeLa cells were seeded at a density of 5 × 10^4^ cells/ml in a 6-well plate. The next day, cells were transfected with 0.2 μg of Super PiggyBac Transposase Expression Vector (System Biosciences, PB210PA-1) and 0.2, 1, or 2 μg of transposon vectors with appropriate “cargo” sub cloned using Lipofectamine 3000 (Invitrogen) according to the manufacturer’s instructions. After 3 days, the cells were treated with 10 μg/ml blasticidin S (Gibco, A1113903) for selection. Cells with positive integrants for more than 7 days were validated using immunoblotting or RT-qPCR following treatment with DOX. Jurkat cells were seeded at a density of 8 × 10^5^ cells/ml in a 6-well plate and transfected with 0.2 µg of transposase and 1 µg of corresponding transposon vectors with Lipofectamine 3000, like HeLa cells. After 3 days, the cells were treated with 10 μg/ml blasticidin S (Gibco, A1113903) for selection. For each passage, cells were cushioned onto Ficoll-Pacque (Cytiva, 17144002) to separate live cells from dead cell debris. The cells over the cushion were washed with PBS and incubated in cell culture medium with 10 µg/ml of blasticidin for further selection for at least 14 days. Cells with positive integrants were validated by immunoblotting after treatment with DOX. Quantification of successfully integrated piggyBac transposons was performed using a piggyBac qPCR copy number kit (System Biosciences, PBC100A-1).

### HIV-1 integration site sequencing library construction

HIV-1 integration site sequencing library construction was performed as described earlier (*7, 9*). Summarily, HeLa cells and primary CD4^+^ T cells were infected with VSV-G-pseudotyped HIV-1 NL4-3 ΔEnv EGFP virus at a MOI of 0.6 and harvested 5 days post infection. gDNA was isolated using a DNA purification kit (Qiagen, 51106), according to the manufacturer’s instructions. gDNA (10 µg) was digested overnight at 37°C with 100 U each of the restriction endonucleases MseI (NEB, R0525L) and BglII (NEB, R0144L). Linker oligonucleotides, which were compatible for ligation with the MseI-generated DNA ends, were ligated with gDNA overnight at 12°C in reactions containing 1.5 μM ligated linker, 1 μg fragmented DNA, and 800 U T4 DNA ligase (NEB, M0202S). Viral LTR–host DNA junctions were amplified using semi-nested PCR with a unique linker-specific primer and LTR primers. The second round of PCR was carried out with primers binding to the LTR and the linkers for next-generation sequencing. Two PCRs were performed in parallel for the first round of PCR and five PCRs were performed in parallel for the second round of PCR to enhance library diversity. S7 Table presents details of the oligonucleotides used for HIV-1 integration site sequencing library construction. HIV-1 integration site sequencing was performed in biological replicates. Integration site libraries were analyzed using 150 bp paired-end sequencing on a HiSeqX Illumina instrument.

### Co-immunoprecipitation of DNA–RNA hybrid

DNA–RNA hybrid immunoprecipitation was performed as described earlier (*39*). Summarily, non-crosslinked HeLa cells transfected with the pFlag-IN codon-optimized plasmid were lysed in 85 mM KCl, 5 mM PIPES (pH 8.0), and 0.5% NP-40 for 10 min on ice, and then, the lysates were centrifuged at 750 *g* for 5 min to pellet the nuclei. The pelleted nuclei were resuspended in sodium deoxycholate, SDS, and sodium lauroyl sarcosinate in RSB buffer and were sonicated for 10 min (Diagenode Bioruptor). Extracts were then diluted (1:4 in RSB + T buffer) and subjected to immunoprecipitation with the S9.6 antibody overnight at 4°C. Antibody-bound complexes were incubated with Protein A Dynabeads (Invitrogen) for 4 h at 4°C for immunoprecipitation. Normal mouse IgG antibodies (Santa Cruz, sc-2025) were used as negative controls. RNase A (Thermo Scientific, EN0531) was added during immunoprecipitation at 0.1 ng RNase A per µg gDNA. Beads were washed four times with RSB + T; twice with RSB, and eluted either in 2× LDS (Novex, NP0007), 100 mM DTT for 10 min at 70°C (for western blot), or 1% SDS and 0.1 M NaHCO_3_ for 30 min at room temperature (for DNA–RNA hybrid dot blot).

For co-immunoprecipitation of DNA–RNA hybrids with RNase H treatment, gDNA containing RNA–DNA hybrids was isolated from HeLa cells transfected with a pFlag-IN codon-optimized plasmid using a QIAmp DNA Mini Kit (Qiagen, 51304). gDNA was sonicated for 10 min (Diagenode Bioruptor) and then treated with 5.5 U RNase H (NEB, M0297) per µg of DNA overnight at 37 °C. A fraction of gDNA was stored as “nucleic acid input” for dot blot analysis. gDNA was cleaned using standard phenol-chloroform extraction, resuspended in DNase/RNase-free water, enriched for DNA–RNA hybrids using immunoprecipitation with the S9.6 antibody (overnight at 4°C), isolated with Protein A Dynabeads (Invitrogen; 4 h at 4°C), washed thrice with RSB+T. The immunoprecipitated complexes were incubated with nuclear extracts of HeLa cells transfected with the pFlag-IN codon-optimized plasmid for 2 h at 4°C with diluted HeLa nuclear extracts. The cell lysate containing proteins were pre-treated with 0.1 mg/ml RNase A (Thermo Scientific, EN0531) for 1 h at 37°C to degrade all RNA–DNA hybrids, and the excess of RNase A was blocked by adding 200 U of SUPERase in RNase inhibitor (Invitrogen, AM2694) for immunoprecipitation. In addition, 100 μL fraction of diluted and RNase A pre-treated extracts prior to immunoprecipitation was stored as “protein input” for western blotting. Beads were washed four times with RSB + T; twice with RSB, and eluted either in 2× LDS (Novex, NP0007), 100 mM DTT for 10 min at 70°C (for western blot), or 1% SDS, and 0.1 M NaHCO_3_ for 30 min at room temperature (for DNA–RNA hybrid dot blot).

### Recombinant Sso7d-IN protein purification

Sso7d-integrase active site mutant E152Q was expressed in Escherichia coli BL21-AI and purified essentially as previously described (*52*). Briefly, Sso7d-IN (E152Q) expressed BL21-AI cells were lysis in lysis buffer (20 mM HEPES pH 7.5, 2 mM 2-mercaptoethanol, 1 M NaCl, 10% (w/v) glycerol, 20 mM imidazole, 1 mg RNase A, and 1000 U DNase I) and purified by nickel affinity chromatography (Qiagen, 30210). Protein were first loaded on HeparinHP column (GE Healthcare) equilibrated with equilibrated with 20 mM Tris, pH 8.0, 0.5 mM TCEP, 200 mM NaCl, 10% glycerol for anion exchange chromatography prior to the size exclusion chromatography. Proteins were eluted with a linear gradient of NaCl from 200 mM to 1 M. Eluted fractions were pooled and then separated on a Superdex-200 PC 10/300 GL column (GE Healthcare) equilibrated with 20 mM Tris pH 8.0, 0.5 mM TCEP, 500 mM NaCl and 6% (w/v) glycerol. The purified protein was concentrated to 0.6 mg/ml using an Amicon centrifugal contentrator (EMD Millipore), flash-frozen in liquid nitrogen and stored at –80°C.

### Electrophoretic mobility shift assay for R-loop binding of Sso7d-IN

To test the binding affinity of Sso7d-tagged HIV-1 IN to different types of nucleic acid substrates, we prepared R-loop, dsDNA, RNA-DNA hybrid with exposed ssDNA (R:D+ssDNA), RNA-DNA hybrid (Hybrid), ssDNA, and ssRNA by annealing different combinations of Cy3, Cy5 or non-labeled oligonucleotides following the previous protocol (*53*). 10 nM of DNA substrate was incubated with Sso7d-IN at different concentrations in assembly buffer (20 mM HEPES pH 7.5, 5 mM CaCl_2_, 8 mM 2-mercaptoethanol, 4 uM ZnCl_2_, 100 mM NaCl, 25% (w/v) glycerol and 50 mM 3-(Benzyldimethylammonio) propanesulfonate (NDSB-256)), for 1 h at 30°C then incubated for 15 min on ice. All the reactants were run on 4.5% non-denaturing PAGE in 1× TBE and then Cy3 or Cy5 fluorescence signal was imaged by ChemiDoc MP imaging system (Bio-Rad). S8 Table presents details of the oligonucleotide sequence used for EMSA.

### PLA

For PLA, HeLa cells were grown on coverslips and infected with HIV-IN-EGFP virions. At 6 hpi, cells were pre-extracted with cold 0.5% NP-40 for 3 min on ice. The cells were fixed with 4% paraformaldehyde in PBS for 15 min at 4 °C. The cells were then blocked with 1× blocking solution (Merck, DUO92102) for 1 h at 37°C in a humidity chamber. After blocking, cells were incubated with the following primary antibodies overnight at 4°C for S9.6-HIV-1-IN_PLA: mouse anti-DNA–RNA hybrid S9.6 (1:250; Kerafast, ENH001) and rabbit anti-GFP (1:500; Abcam, ab6556). The following day, after washing with once with buffer A twice (Merck, DUO92102), cells were incubated with pre-mixed Duolink PLA plus (anti-mouse) and PLA minus probes (anti-rabbit) antibodies for 1 h at 37°C. The subsequent steps in the proximal ligation assay were performed using the Duolink PLA Fluorescence kit (Sigma) according to the manufacturer’s instructions. To obtain images, the mounted specimens were visually scanned and representative images were acquired using a Zeiss LSM 710 laser scanning confocal microscope (Carl Zeiss). The number of intranuclear PLA puncta was quantified using the ImageJ software. For each biological replicate and experiment, a PLA with a single antibody was performed as a negative control under the same conditions.

### DRIPc-Seq data processing and peak calling

DRIPc-seq reads were quality-controlled using FastQC v0.11.9 (*54*), and sequencing adapters were trimmed using Trim Galore! v0.6.6 (*55*) based on Cutadapt v2.8 (*56*). Trimmed reads were aligned to the hg38 reference genome using bwa v0.7.17-r1188 (*57*). Read deduplication and peak calling were performed using MACS v2.2.7.1 (*58*). Because R-loops appear as both narrow and broad peaks in DRIPc-seq read alignment owing to its variable length, two independent “MACS2 callpeak” runs were performed for narrow and broad peak calling. The narrow and broad peaks were merged using Bedtools v2.26.0 (*59*). To increase the sensitivity of DRIPc-seq peak identification, peaks were called after pooling the two biological replicates of the DRIPc-seq sequencing data for each condition.

### Consensus R-loop peak calling

The R-loop peaks at 0, 3, 6 and 12 hpi were first merged using “bedtools merge” to create a universal set of R-loop peaks across time points (n = 46542). Then, each of the universal R-loop peaks was tested for overlap with the R-loop peaks for 0, 3, 6 and 12 hpi using “bedtools intersect”. In all, 9,190, 21,403, 33,544, and 9,941 peaks overlapped with 0, 3, 6, and 12 hpi R-loop peaks, respectively. For CD4 cells, we identified a universal R-loop set consisting of 3,928 R-loops, and among them, 737, 722, 1,796 and 2,766 peaks overlapped with 0, 3, 6 and 12hpi R-loop peaks.

### HIV-1 integration site sequencing data processing

Quality control of HIV-1 integration site-sequencing reads was performed using FastQC v0.11.9. To discard primers and linkers specific for integration site-sequencing from reads, we used Cutadapt v2.8 with the following option: “-u 49 –U 38 –-minimum-length 36 –-pair-filter any –-action trim –q0,0 –a linker –A TGCTAGAGATTTTCCACACTGACTGGGTCTGAGGG –A GGGTCTGAGGG –-no-indels –-overlap 12”. This allowed the first position of the read alignment to directly represent the genomic position of HIV-1 integration. Processed reads were aligned to the hg38 reference genome using bwa v0.7.17-r1188, and integration sites were identified using an in-house Python script. Genomic positions supported by more than five read alignments were regarded as HIV-1 integration sites. For Jurkat cells, we adopted integration site sequencing data of HIV-1 infected wild type Jurkat cells from SRR12322252 (*60*).

### Co-localization analysis of R-loops and integration sites

Enrichment of integration sites near the R-loop peaks was tested using a randomized permutation test. Randomized R-loop peaks were generated using “bedtools shuffle” command, thus preserving the number and the length distribution of the R-loop peaks during the randomization process. Similarly, integration sites were randomized using the “bedtools shuffle” command. Randomization was performed 100 times. ENCODE blacklist regions (*61*) were excluded while shuffling the R-loops and integration sites to exclude inaccessible genomic regions from the analysis. For each of the observed (or randomized) integration sites, the closest observed (or randomized) R-loop peak and the corresponding genomic distance were identified using the “bedtools closest” command. The distribution of the genomic distances was displayed to show the local enrichment of integration sites in terms of the increased proportion of integration sites within the 30-kb window centered on R-loops compared to their randomized counterparts.

### DNA plasmid construction and transfection

R-loop-forming mAIRN and non-R-loop forming ECPF sequences were subcloned from pSH26 and pSH36 plasmids, which were generously provided by Prof. Karlene A. Cimprich, into the piggyBac transposon vector, where the tet operator sequences were located upstream of the minimal CMV promoter. The pFlag-IN codon-optimized plasmid and pVpr-IN-EGFP were kindly provided by Prof. A. Engelman and Prof. Anna Cereseto, respectively. Lipofectamine 3000 (Invitrogen) transfection reagent was used for the transfection of all plasmids into cells, according to the manufacturer’s protocol.

### DNA–RNA hybrid dot blotting

Total gDNA was extracted using the QIAmp DNA Mini Kit (Qiagen, 51304) according to the manufacturer’s instructions. gDNA (1.2 μg) was treated with 2 U RNase H (NEB, M2097) per µg of gDNA for 4 h at 37°C, with half of the sample left untreated but denatured. Half of the DNA sample was probed with S9.6 antibody (1:1000), and the other half was probed with an anti-ssDNA antibody (MAB3034, Millipore, 1:10000).

### Immunoblotting

Cells were lysed using RIPA buffer (50 mM Tris, 150 mM sodium chloride, 0.5% sodium deoxycholate, 0.1% SDS, and 1.0% NP-40) supplemented with 10 μM leupeptin (Sigma-Aldrich) and 1 mM phenylmethanesulfonyl fluoride (Sigma-Aldrich) and boiled at 98°C for 10 min with SDS sample buffer prior to SDS-PAGE. The primary antibodies used were mouse monoclonal anti-FLAG M2 (Sigma, F3165), monoclonal mouse anti-HSC70 (Abcam, ab2788), polyclonal rabbit anti-histone H3 (tri methyl K4) antibody (Abcam, ab8580), monoclonal mouse anti-HIV-1 Integrase (Santa Cruz, sc-69721), rabbit anti-LaminA/C antibody (Cell Signaling, 2032), and monoclonal mouse anti-Actin (Invitrogen, MA1-744). All primary antibodies were used at a dilution of 1:1000 for western blotting. Peroxidase-conjugated anti-mouse IgG (115-035-062) and anti-rabbit IgG (111-035-003; both Jackson Laboratories) were used as secondary antibodies at 1:5000 dilution. Signals were detected using the SuperSignal West Pico chemiluminescence kit (Thermo Fisher Scientific).

### RNA-seq data processing

RNA-seq reads were quality-controlled and adapter-trimmed as in DRIPc-seq processing. To quantify the expression levels of protein-coding genes, processed reads were aligned to the hg38 reference genome with GENCODE v37 gene annotation (*62*) using STAR v2.7.3a (*63*). Gene expression quantification was performed using RSEM v1.3.1. To quantify the expression levels of transposable elements (TEs), we used TEtranscripts v2.2.1 (*64*). Processed reads were first aligned to the hg38 reference genome using GENCODE v37 and RepeatMasker TE annotation using STAR v2.7.3a. In this case, STAR options were modified as follows to utilize multimapping reads in downstream analyses: “--outFilterMultimapNmax 100 –-winAnchorMultimapNmax 100 –-outMultimapperOrder random –-runRNGseed 77 –-outSAMmultNmax 1 –-outFilterType BySJout –-alignSJoverhangMin 8 –-alignSJDBoverhangMin 1 –-alignIntronMin 20 –-alignIntronMax 1000000 –-alignMatesGapMax 1000000”. Expression levels of TEs were quantified as read counts with the “TEcount” command.

### Genome annotations

All bioinformatic analyses were performed using the hg38 reference genome and GENCODE v37 gene annotation. Promoters were defined as a 2-kb region centered at the transcription start sites of the APPRIS principal isoform of protein-coding genes. TTS regions were defined as the 2-kb region centered at the 3′ terminals of protein-coding transcripts. CpG island annotations were downloaded from the UCSC table browser. CpG shores were defined as 2-kb regions flanking CpG islands, excluding the regions overlapping with CpG islands. Similarly, CpG shelves were defined as 2-kb regions flanking the stretch of CpG islands and shores while excluding the regions overlapping with CpG islands and shores. Annotations for LINE, SINE, and LTR were extracted from the RepeatMasker track in the UCSC table browser.

### Identification of viral sequencing reads in DRIPc-seq

To identify sequencing reads originating from the viral genome, we aligned DRIPc-seq reads to a composite reference genome consisting of the human and HIV1 genome (Genbank accession number: AF324493.2) and computed the proportion of the reads mapped to HIV1 genome.

## Code availability

Bioinformatics pipelines and scripts used in this study are accessible from https://github.com/dohlee/hiv1-rloop.

## Acknowledgements

We are grateful to Prof. Karlene A. Cimprich (Standford University) for providing the pSH26 and pSH36 plasmids, Prof. A. Engelman (Harvard Medical School) for providing pFlag-IN codon optimized plasmid and Prof. Anna Cereseto (University of Trento) for providing pVpr-IN-EGFP. The NL4-3 ΔEnv EGFP and pNL4-3.Luc.R-E-viral plasmids were obtained through the NIH HIV Reagent Program, Division of AIDS, NIAID, NIH. We thank Dr. Sungchul Kim (IBS center for RNA Research) and Seongjin An (Korea University) for their technical support in recombinant protein purification.

## Author contributions

K.P. and K.A. designed experiments. K.P., J.J and S.L. performed experiments. D.L. performed the bioinformatical and statistical analyses. K.P., D.L., K.A. and S.K. analyzed the data. K.P., D.L., and K.A. wrote the manuscript.

## Supplemental Information

### Materials and Methods

S1 Fig. Primary CD4^+^ T cells sorting strategies and GFP-HIV-1 infection

S2 Fig. Genome browser screenshot over the HIV-1-induced R-loop forming positive or negative genomic regions

S3 Fig. Host cellular R-loop induction by HIV-1 infection is host-genome specific.

S4 Fig. R-loop induction by HIV-1 infection does not follow transcriptome changes in HeLa cells

S5 Fig. PiggyBac transposon-transposase insertion of R-loop forming and non-R-loop forming sequences in HeLa cells

S6 Fig. HIV-1 integrase proteins directly binds to host genomic R-loops

S1 Table. Chromosomal position and DRIPc-seq signal for referenced R-loop-positive and – negative regions in HIV-1 infected HeLa cells

S2 Table. Chromosomal position and DRIPc-seq signal for referenced R-loop-positive and – negative regions in HIV-1 infected primary CD4^+^ T cells

S3 Table. Chromosomal position and DRIPc-seq signal for referenced R-loop-positive and – negative regions in HIV-1 infected Jurkat cells

S4 Table. RNA-seq analysis of relative gene expression levels of P1-3 and N1,2 R-loop regions

S5 Table. Oligonucleotides used for DRIPc-seq library construction

S6 Table. Primers used for qPCR

S7 Table. Oligonucleotides used for HIV-1 integration site sequencing library construct

S8 Table. Oligonucleotides used for electrophoretic mobility shift assay substrate preparation

S9 Table. Accession numbers and data sources.

### Supplementary figures

**S1 Fig.**
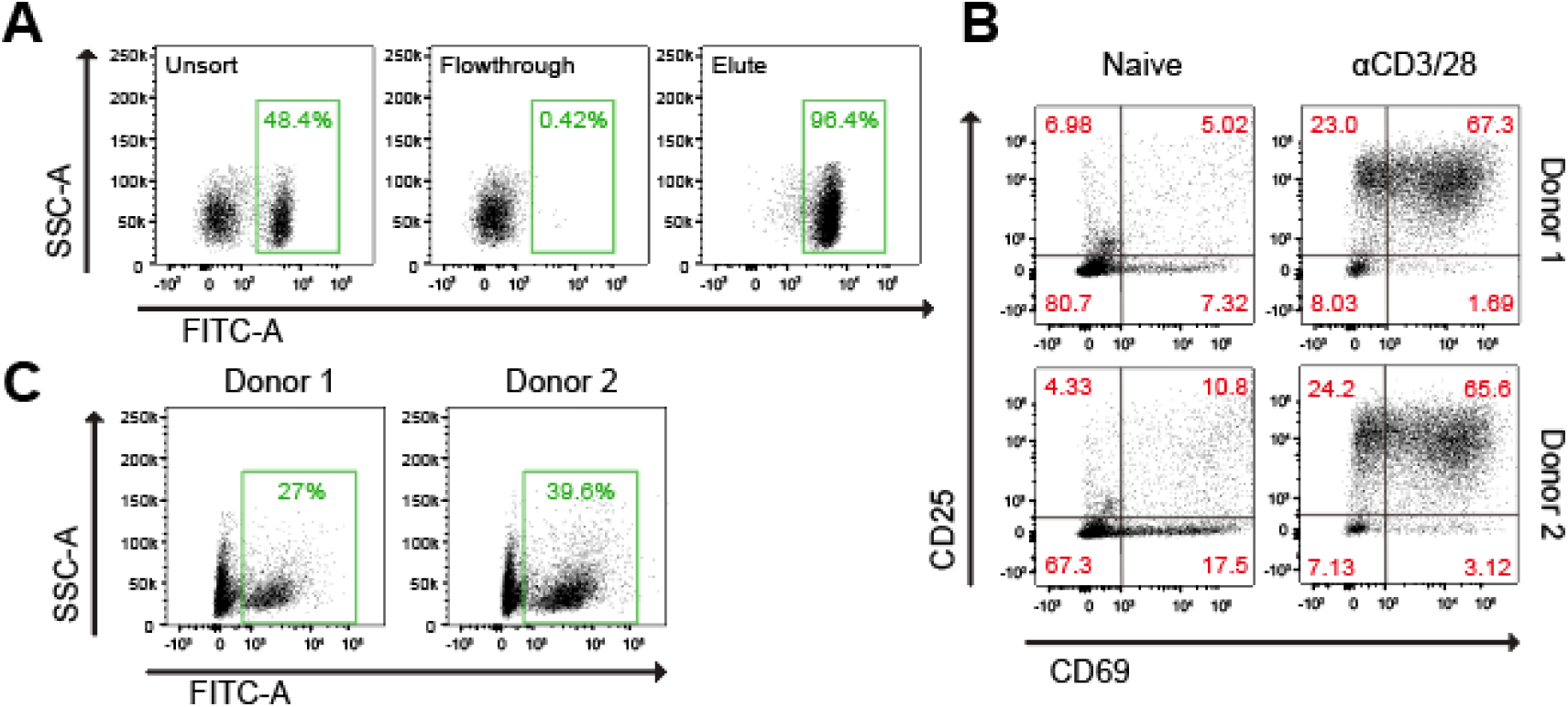
Primary CD4^+^ T cells sorting strategies and GFP-HIV-1 infection. (**A**) Gating strategy used to determine the efficiency of CD4^+^ T cells sorting from human PBMC. Pre-sorted PBMCs were staining with FITC-conjugated anti-CD4 and subjected for positive CD4+ T cell sorting. The percentages of FITC stained cell population at each step of cell sorting are as indicated. (**B**) Gating strategy used to determine non-activated (Naïve) and activated cells (αCD3/28) with two markers, CD25 (FITC) and CD69 (APC), for each donor (upper panels, Donor 1; lower panels, Donor 2). (**C**) Gating strategy used to determine HIV-1-infectivity of CD4+ T cells from each donor infected with GFP reporter HIV-1 virus at 48 hpi. The percentages of GFP positive cell population at are as indicated.

**S2 Fig.**
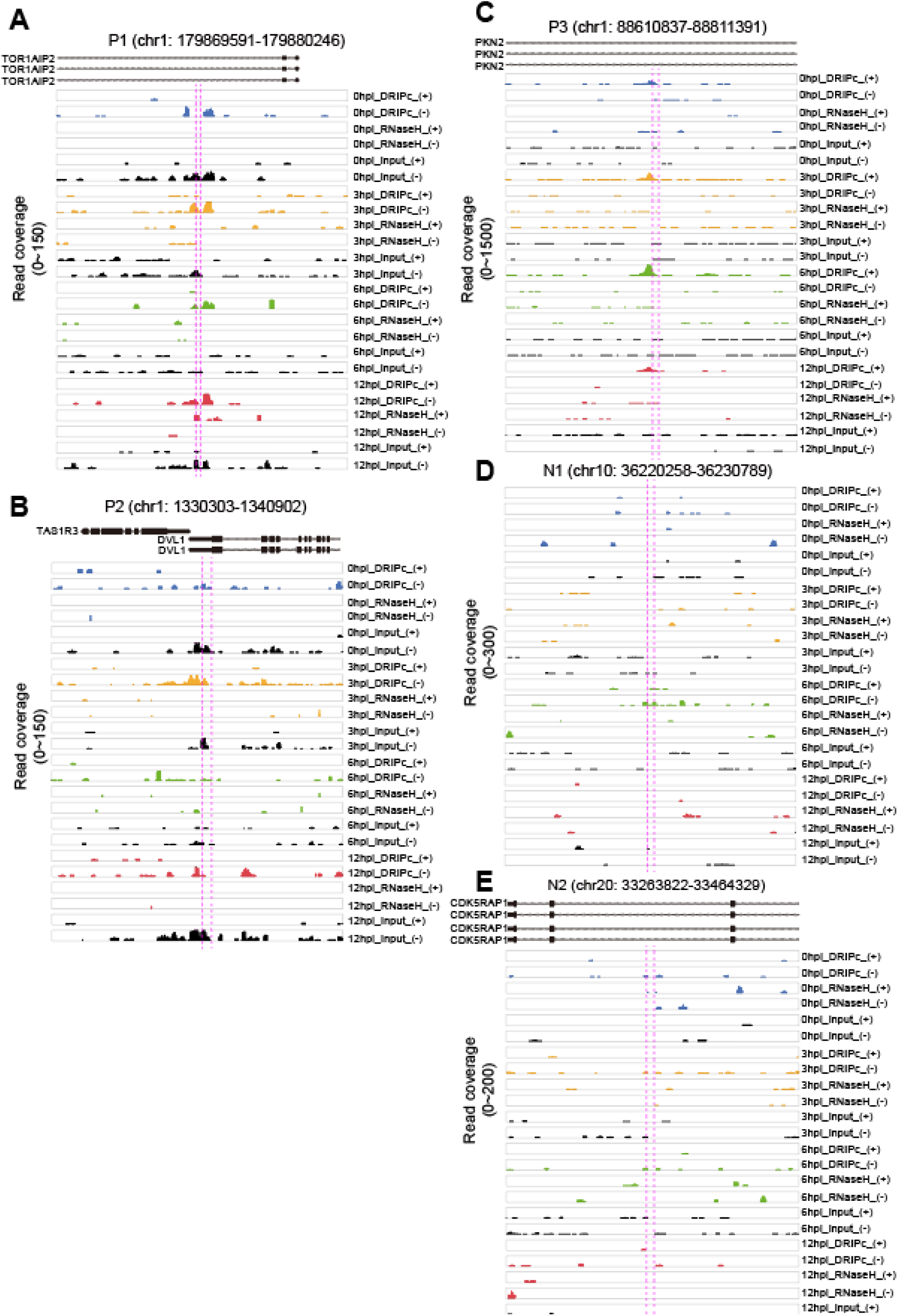
Genome browser screenshot over the HIV-1-induced R-loop forming positive or negative genomic regions. (**A-C**), Genome browser screenshot over the P1 (**A**), P2 (**B**), and P3 (**C**) HIV-1 induced R-loop-positive chromosomal regions showing result from DRIPc-seq in HIV-1-infected HeLa cells (blue, 0 hpi; yellow, 3 hpi; green, 6 hpi; red, 12 hpi; black, input signals for each indicated sample) on plus (+) or minus (-) DNA strand. Magenta dotted lines represent primer binding sites in qPCR following DRIP. **(D** and **E),** Genome browser screenshot over the N1 (**D**), and N2 (**E**) HIV-1 induced R-loop-negative chromosomal regions showing result from DRIPc-seq in HIV-1-infected HeLa cells (blue, 0 hpi; yellow, 3 hpi; green, 6 hpi; red, 12 hpi; black, input signals for each indicated sample) on plus (+) or minus (-) DNA strand. Magenta dotted lines represent primer binding sites in qPCR following DRIP.

**S3 Fig.**
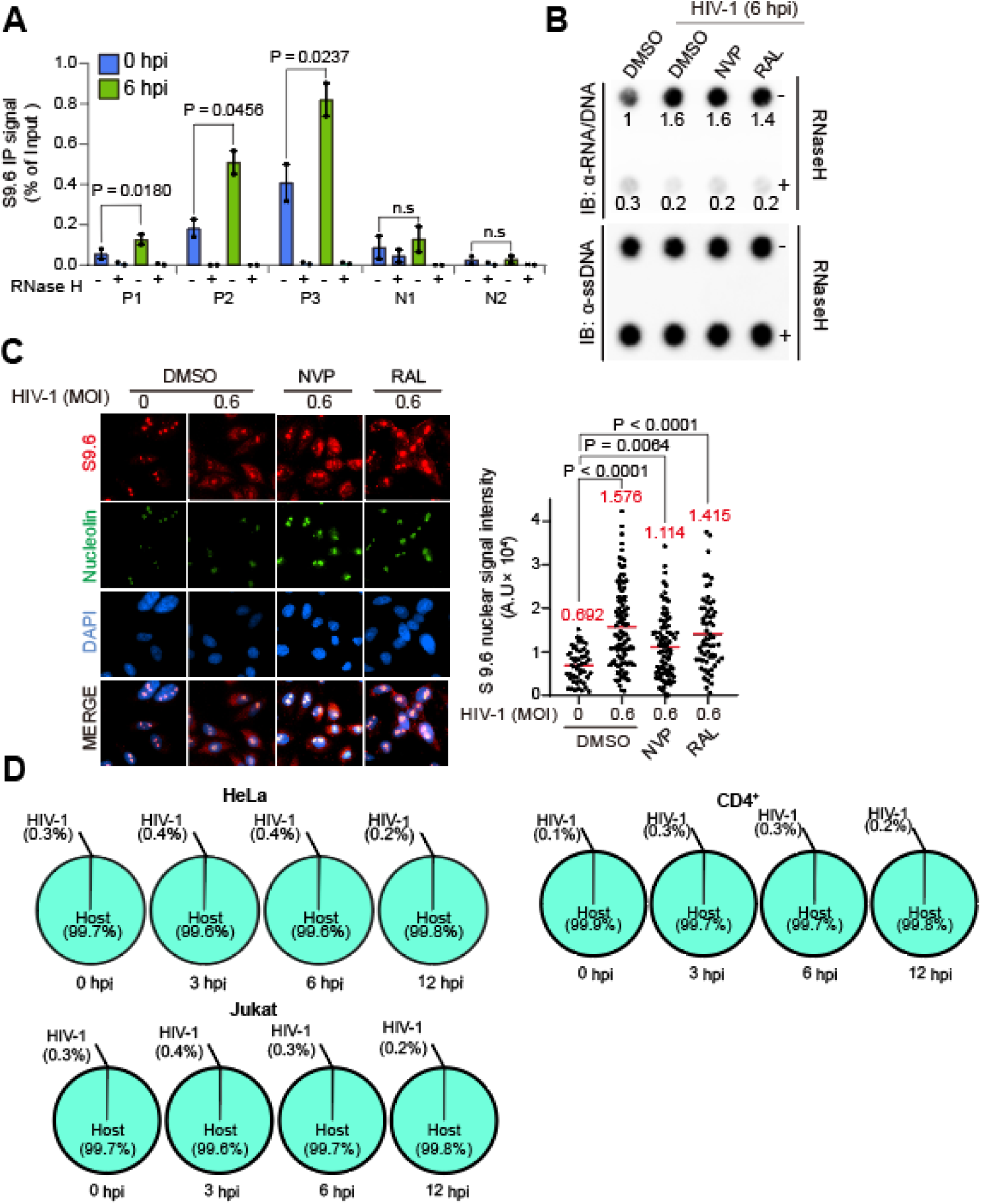
Host cellular R-loop induction by HIV-1 infection is host-genome specific. (**A**) DRIP-qPCR using the anti-S9.6 antibody at P1, P2, P3, N1, and N2 in HIV-1-infected cells with MOI of 0.6 harvested at the indicated hpi (blue, 0 hpi; green, 6 hpi). Pre-immunoprecipitated materials were untreated (−) or treated (+) with RNase H, as indicated. Data are presented as the mean ± SEM; P-values were calculated using one-way ANOVA (n = 2). (**B**) Dot blot analysis of the R-loop in gDNA extracts from HIV-1 infected HeLa cells with MOI of 0.6 harvested at 6hpi. The cells were treated with DMSO, 10uM of Nevirapine (NVP), or 10uM of Raltegravir (RAL) for 24 h before infection, as indicated. gDNAs were probed with anti-S9.6. gDNA extracts were incubated with or without RNase H in vitro before membrane loading (anti-RNA/DNA signal). Fold-induction was normalized to the value of harvested cells at 0 hpi by quantifying the dot intensity of the blots and calculating the ratios of the S9.6 signal to the total amount of gDNA (anti-ssDNA signal). (**C**) Representative images of the immunofluorescence assay of S9.6 nuclear signals in HIV-1 infected HeLa cells with MOI of 0.6 at 6 hpi. The cells were pre-extracted of cytoplasm and co-stained with anti-S9.6 (red), anti-nucleolin antibodies (green), and DAPI (blue). The cells were treated with DMSO, 10uM of Nevirapine (NVP), or 10uM of Raltegravir (RAL) for 24 h before infection, as indicated. Quantification of S9.6 signal intensity per nucleus after nucleolar signal subtraction for the immunofluorescence assay. The mean value for each data point is indicated by the red line. Statistical significance was assessed using one-way ANOVA (n >51). (**D**) Pie graphs indicating the percentage of DRIPc-seq reads aligned to host cellular genome (aquamarine) or to HIV-1 viral genome (gray), out of the total consensus DRIPc-seq peaks from HIV-infected HeLa cells, primary CD4^+^ T cells and Jurkat cells.

**S4 Fig.**
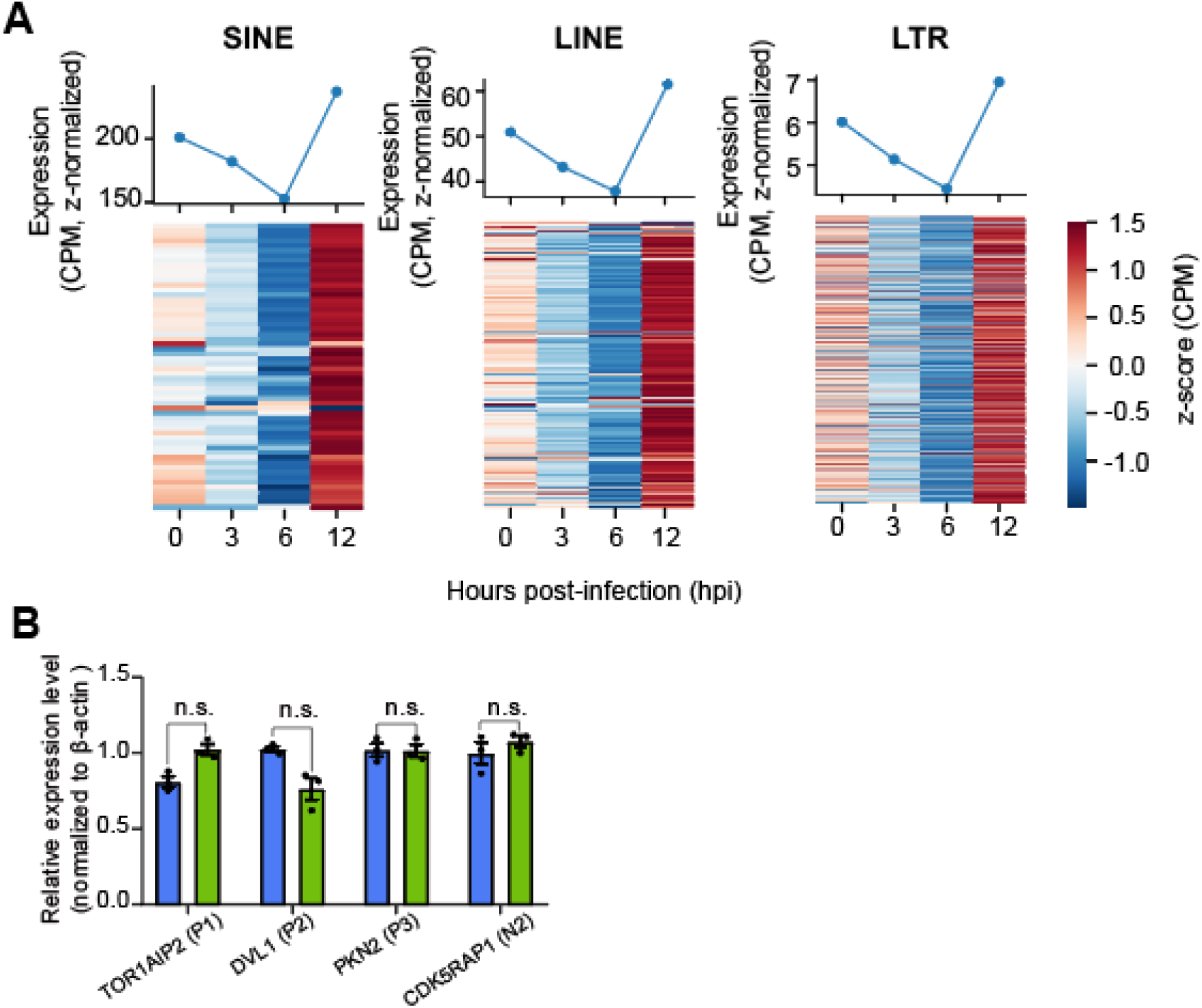
R-loop induction by HIV-1 infection does not follow transcriptome changes in HeLa cells. (**A**) Line graphs and heat maps representing expression levels of indicated repetitive elements (SINE, right; LINE, middle; LTR, left) at the indicated hpi of HIV-1 in HeLa cells. Data are presented as the mean expression levels of two biologically independent experiments. (**B**) Indicated gene expression as measured by RT-qPCR in 0 or 6 hpi harvested HIV-1-infected HeLa cells. Data represent mean ± SEM, n = 3, P values were calculated according to two-tailed Student’s t-test. P > 0.05; n.s, not significant.

**S5 Fig.**
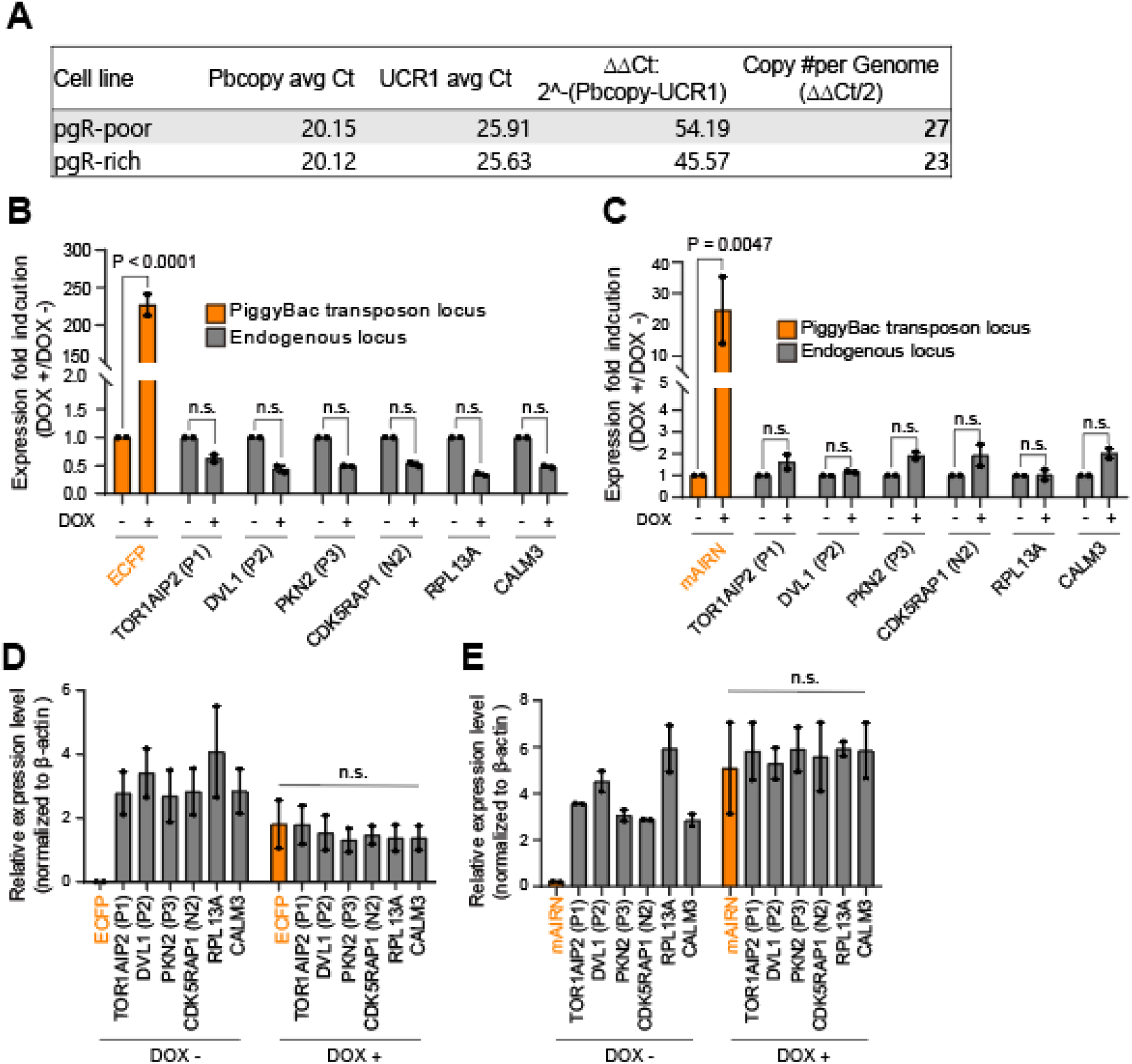
Regulation of cellular R-loops by RNase H1 expression, or by transposon-transposase insertion of R-loop forming and non-R-loop forming sequences in HeLa cells. (**A**) Copy number of piggyBac transposon inserts in each cell line constructed by transfecting the transposon vector and transposase-expressing vector. (**B** and **C**) Fold induction of gene expression for the indicated genes as measured by RT-qPCR. Fold induction were calculated by dividing the gene expression level of DOX-treated (+) by that of DOX-untreated (-) in pgR-poor cells (**B**) or pgR-rich cells (**C**). Data represent mean ± SEM, n = 2, P values were calculated according to two-way ANOVA. P > 0.05; n.s, not significant. (**D** and **E**) Relative gene expression of the indicated genes as measured by RT-qPCR in DOX-treated (+) or DOX-untreated (-) pgR-poor cells (**D**) or pgR-rich cells (**E**). Data represent mean ± SEM, n = 2, P values were calculated according to two-way ANOVA. P > 0.05; n.s, not significant.

**S6 Fig.**
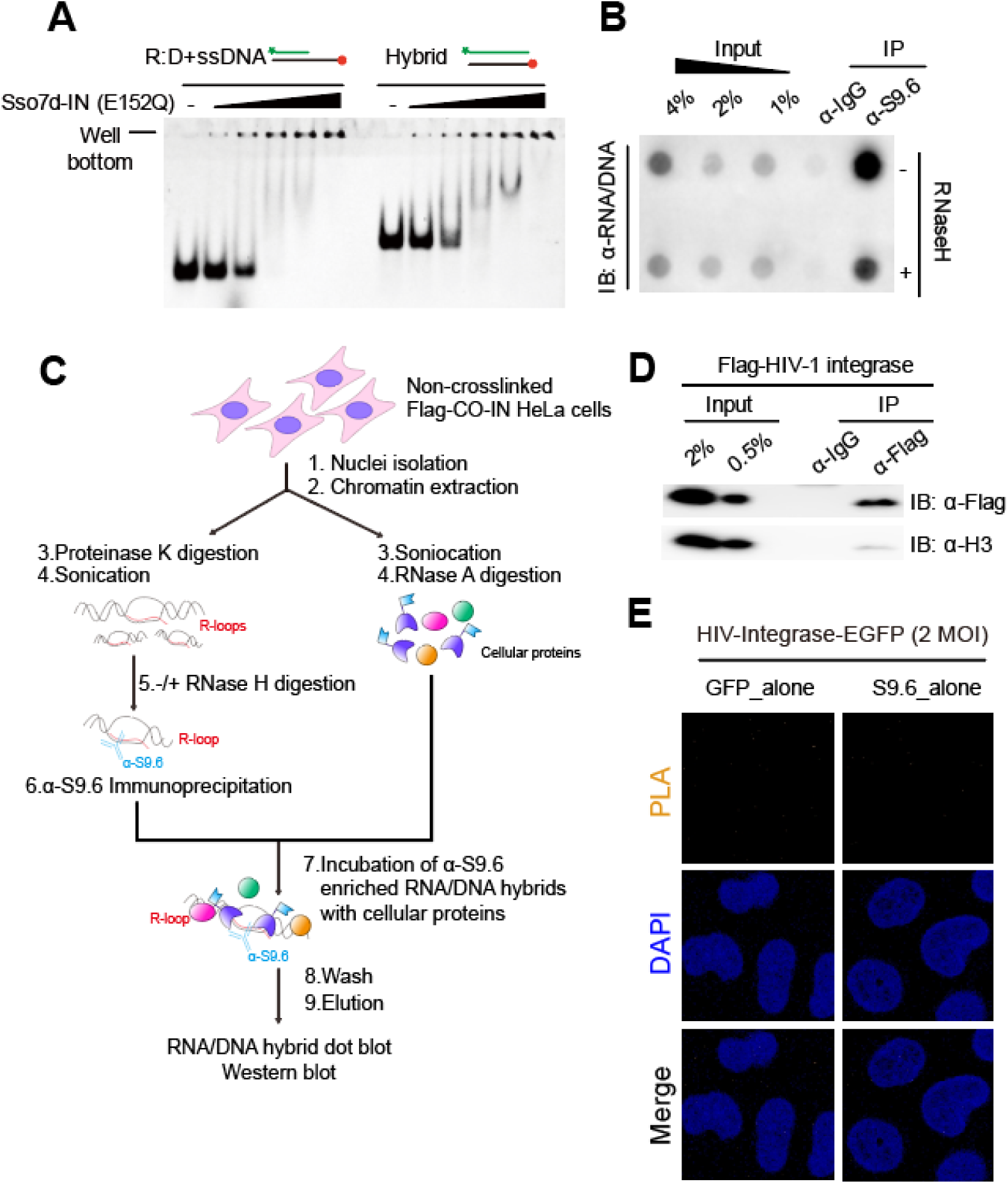
HIV-1 integrase proteins directly binds to host genomic R-loops. (**A**) Representative gel images for EMSA of Sso7d-tagged HIV-1-integrase (E152Q) with different types of nucleic acids substrates (R:D+ssDNA and Hybrid). 100 nM nucleic acid substrate was incubated with Sso7d-tagged HIV-1-integrase (E152Q) at 0 nM, 20 nM, 50 nM, 100 nM, 200 nM, and 400 nM (n = 3). (**B**) Nucleic acid extracts from FLAG-HIV-1-integrase-expressing cells were immunoprecipitated using S9.6 antibody. gDNA was precipitated from the elutes of immunoprecipitation and subjected to DNA–RNA hybrid dot blotting. Where indicated, the gDNA extracts were either untreated (−) or treated (+) with RNase H after elution of immunoprecipitated materials. (**C**) Summary of the experimental design for R-loop immunoprecipitation using S9.6 antibody in FLAG-tagged HIV-1 integrase protein-expressing HeLa cells with pre-immunoprecipitation in vitro RNase H treatment. (**D**) Protein extracts from FLAG-HIV-1-integrase-expressing cells were immunoprecipitated using anti-FLAG antibody. Western blot of FLAG immunoprecipitation was probed with anti-FLAG or anti-H3 antibodies. (**E**) Representative images of the proximity-ligation assay (PLA) using single antibody (anti-GFP or anti-S9.6) in HIV-IN-EGFP virion-infected HeLa cells at 6 hpi, as PLA signal negative controls. Cells were subjected to PLA (orange) and co-stained with DAPI (blue) (n > 50).

**S1 Table.** Chromosomal position and DRIPc-seq signal for referenced R-loop-positive and –negative regions in HIV-1 infected HeLa cells.

| HeLa |  |  |  |  |
| --- | --- | --- | --- | --- |
| Gene | Chromosom | Position (hg38) | Description | Average DRIPc-seq signal |
| RPL13A | chr19 | 49487608-49493057 | Input (-)_0hpi | 3.59 |
|  |  |  | Input (-)_3hpi | 0.24 |
|  |  |  | Input (-)_6hpi | 2.39 |
|  |  |  | Input (-)_12hpi | 3.51 |
|  |  |  | Input (+)_0hpi | 82.29 |
|  |  |  | Input (+)_3hpi | 51.76 |
|  |  |  | Input (+)_6hpi | 39.14 |
|  |  |  | Input (+)_12hpi | 176.73 |
|  |  |  | IP_RNase H- (-)_0hpi | 2.21 |
|  |  |  | IP_RNase H- (-)_3hpi | 2.73 |
|  |  |  | IP_RNase H- (-)_6hpi | 2.39 |
|  |  |  | IP_RNase H- (-)_12hpi | 4.25 |
|  |  |  | IP_RNase H- (+)_0hpi | 110.32 |
|  |  |  | IP_RNase H- (+)_3hpi | 140.22 |
|  |  |  | IP_RNase H- (+)_6hpi | 58.36 |
|  |  |  | IP_RNase H- (+)_12hpi | 137.37 |
|  |  |  | IP_RNase H+ (-)_0hpi | 0.00 |
|  |  |  | IP_RNase H+ (-)_3hpi | 4.48 |
|  |  |  | IP_RNase H+ (-)_6hpi | 3.74 |
|  |  |  | IP_RNase H+ (-)_12hpi | 0.00 |
|  |  |  | IP_RNase H+ (+)_0hpi | 1.98 |
|  |  |  | IP_RNase H+ (+)_3hpi | 3.36 |
|  |  |  | IP_RNase H+ (+)_6hpi | 1.60 |
|  |  |  | IP_RNase H+ (+)_12hpi | 6.81 |
| Gene | Chromosom | Position (hg38) | Description | Average DRIPc-seq signal |
| CALM3 | chr19 | 46601330-46610782 | Input (-)_0hpi | 1.47 |
|  |  |  | Input (-)_3hpi | 1.02 |
|  |  |  | Input (-)_6hpi | 2.46 |
|  |  |  | Input (-)_12hpi | 0.74 |
|  |  |  | Input (+)_0hpi | 26.50 |
|  |  |  | Input (+)_3hpi | 19.95 |
|  |  |  | Input (+)_6hpi | 11.61 |
|  |  |  | Input (+)_12hpi | 56.92 |
|  |  |  | IP_RNase H- (-)_0hpi | 0.90 |
|  |  |  | IP_RNase H- (-)_3hpi | 1.54 |
|  |  |  | IP_RNase H- (-)_6hpi | 1.23 |
|  |  |  | IP_RNase H- (-)_12hpi | 1.73 |
|  |  |  | IP_RNase H- (+)_0hpi | 13.97 |
|  |  |  | IP_RNase H- (+)_3hpi | 28.68 |
|  |  |  | IP_RNase H- (+)_6hpi | 10.58 |
|  |  |  | IP_RNase H- (+)_12hpi | 24.70 |
|  |  |  | IP_RNase H+ (-)_0hpi | 0.71 |
|  |  |  | IP_RNase H+ (-)_3hpi | 1.83 |
|  |  |  | IP_RNase H+ (-)_6hpi | 2.78 |
|  |  |  | IP_RNase H+ (-)_12hpi | 1.04 |
|  |  |  | IP_RNase H+ (+)_0hpi | 2.12 |
|  |  |  | IP_RNase H+ (+)_3hpi | 1.64 |
|  |  |  | IP_RNase H+ (+)_6hpi | 2.26 |
|  |  |  | IP_RNase H+ (+)_12hpi | 1.65 |
| Gene | Chromosom | Position (hg38) | Description | Average DRIPc-seq signal |
| SNRPN | chr15 | 24823647-24978582 | Input (-)_0hpi | 1.46 |
|  |  |  | Input (-)_3hpi | 1.27 |
|  |  |  | Input (-)_6hpi | 1.34 |
|  |  |  | Input (-)_12hpi | 1.76 |
|  |  |  | Input (+)_0hpi | 1.21 |
|  |  |  | Input (+)_3hpi | 0.81 |
|  |  |  | Input (+)_6hpi | 1.25 |
|  |  |  | Input (+)_12hpi | 0.41 |
|  |  |  | IP_RNase H- (-)_0hpi | 0.45 |
|  |  |  | IP_RNase H- (-)_3hpi | 0.47 |
|  |  |  | IP_RNase H- (-)_6hpi | 0.37 |
|  |  |  | IP_RNase H- (-)_12hpi | 0.05 |
|  |  |  | IP_RNase H- (+)_0hpi | 0.37 |
|  |  |  | IP_RNase H- (+)_3hpi | 0.24 |
|  |  |  | IP_RNase H- (+)_6hpi | 0.54 |
|  |  |  | IP_RNase H- (+)_12hpi | 0.07 |
|  |  |  | IP_RNase H+ (-)_0hpi | 1.40 |
|  |  |  | IP_RNase H+ (-)_3hpi | 0.93 |
|  |  |  | IP_RNase H+ (-)_6hpi | 1.10 |
|  |  |  | IP_RNase H+ (-)_12hpi | 1.31 |
|  |  |  | IP_RNase H+ (+)_0hpi | 1.18 |
|  |  |  | IP_RNase H+ (+)_3hpi | 1.12 |
|  |  |  | IP_RNase H+ (+)_6hpi | 1.26 |
|  |  |  | IP_RNase H+ (+)_12hpi | 1.10 |

**S2 Table.** Chromosomal position and DRIPc-seq signal for referenced R-loop-positive and –negative regions in HIV-1 infected primary CD4^+^ T cells.

| CD4+ |  |  |  |  |
| --- | --- | --- | --- | --- |
| Gene | Chromosom | Position (hg38) | Description | Average DRIPc-seq signal |
| RPL13A | chr19 | 49487608-49493057 | Input (-)_0hpi | 2.33 |
|  |  |  | Input (-)_3hpi | 1.51 |
|  |  |  | Input (-)_6hpi | 2.56 |
|  |  |  | Input (-)_12hpi | 0.77 |
|  |  |  | Input (+)_0hpi | 2.91 |
|  |  |  | Input (+)_3hpi | 1.94 |
|  |  |  | Input (+)_6hpi | 2.36 |
|  |  |  | Input (+)_12hpi | 2.19 |
|  |  |  | IP_RNase H- (-)_0hpi | 0.00 |
|  |  |  | IP_RNase H- (-)_3hpi | 3.63 |
|  |  |  | IP_RNase H- (-)_6hpi | 0.00 |
|  |  |  | IP_RNase H- (-)_12hpi | 0.00 |
|  |  |  | IP_RNase H- (+)_0hpi | 144.19 |
|  |  |  | IP_RNase H- (+)_3hpi | 77.26 |
|  |  |  | IP_RNase H- (+)_6hpi | 130.86 |
|  |  |  | IP_RNase H- (+)_12hpi | 190.08 |
|  |  |  | IP_RNase H+ (-)_0hpi | 1.42 |
|  |  |  | IP_RNase H+ (-)_3hpi | 0.00 |
|  |  |  | IP_RNase H+ (-)_6hpi | 0.00 |
|  |  |  | IP_RNase H+ (-)_12hpi | 0.00 |
|  |  |  | IP_RNase H+ (+)_0hpi | 0.93 |
|  |  |  | IP_RNase H+ (+)_3hpi | 0.00 |
|  |  |  | IP_RNase H+ (+)_6hpi | 0.00 |
|  |  |  | IP_RNase H+ (+)_12hpi | 2.28 |
| Gene | Chromosom | Position (hg38) | Description | Average DRIPc-seq signal |
| CALM3 | chr19 | 46601330-46610782 | Input (-)_0hpi | 4.58 |
|  |  |  | Input (-)_3hpi | 4.64 |
|  |  |  | Input (-)_6hpi | 2.96 |
|  |  |  | Input (-)_12hpi | 4.04 |
|  |  |  | Input (+)_0hpi | 3.62 |
|  |  |  | Input (+)_3hpi | 3.65 |
|  |  |  | Input (+)_6hpi | 3.40 |
|  |  |  | Input (+)_12hpi | 4.11 |
|  |  |  | IP_RNase H- (-)_0hpi | 0.00 |
|  |  |  | IP_RNase H- (-)_3hpi | 0.00 |
|  |  |  | IP_RNase H- (-)_6hpi | 0.00 |
|  |  |  | IP_RNase H- (-)_12hpi | 2.70 |
|  |  |  | IP_RNase H- (+)_0hpi | 108.23 |
|  |  |  | IP_RNase H- (+)_3hpi | 183.80 |
|  |  |  | IP_RNase H- (+)_6hpi | 87.73 |
|  |  |  | IP_RNase H- (+)_12hpi | 181.80 |
|  |  |  | IP_RNase H+ (-)_0hpi | 2.80 |
|  |  |  | IP_RNase H+ (-)_3hpi | 0.00 |
|  |  |  | IP_RNase H+ (-)_6hpi | 0.00 |
|  |  |  | IP_RNase H+ (-)_12hpi | 1.94 |
|  |  |  | IP_RNase H+ (+)_0hpi | 4.11 |
|  |  |  | IP_RNase H+ (+)_3hpi | 1.19 |
|  |  |  | IP_RNase H+ (+)_6hpi | 9.88 |
|  |  |  | IP_RNase H+ (+)_12hpi | 6.17 |
| Gene | Chromosom | Position (hg38) | Description | Average DRIPc-seq signal |
| SNRPN | chr15 | 24823647-24978582 | Input (-)_0hpi | 1.65 |
|  |  |  | Input (-)_3hpi | 1.41 |
|  |  |  | Input (-)_6hpi | 1.74 |
|  |  |  | Input (-)_12hpi | 1.15 |
|  |  |  | Input (+)_0hpi | 1.72 |
|  |  |  | Input (+)_3hpi | 1.46 |
|  |  |  | Input (+)_6hpi | 1.97 |
|  |  |  | Input (+)_12hpi | 1.29 |
|  |  |  | IP_RNase H- (-)_0hpi | 0.31 |
|  |  |  | IP_RNase H- (-)_3hpi | 0.27 |
|  |  |  | IP_RNase H- (-)_6hpi | 0.10 |
|  |  |  | IP_RNase H- (-)_12hpi | 0.27 |
|  |  |  | IP_RNase H- (+)_0hpi | 0.98 |
|  |  |  | IP_RNase H- (+)_3hpi | 1.00 |
|  |  |  | IP_RNase H- (+)_6hpi | 0.53 |
|  |  |  | IP_RNase H- (+)_12hpi | 0.56 |
|  |  |  | IP_RNase H+ (-)_0hpi | 0.94 |
|  |  |  | IP_RNase H+ (-)_3hpi | 1.57 |
|  |  |  | IP_RNase H+ (-)_6hpi | 0.00 |
|  |  |  | IP_RNase H+ (-)_12hpi | 2.17 |
|  |  |  | IP_RNase H+ (+)_0hpi | 1.37 |
|  |  |  | IP_RNase H+ (+)_3hpi | 1.14 |
|  |  |  | IP_RNase H+ (+)_6hpi | 1.42 |
|  |  |  | IP_RNase H+ (+)_12hpi | 1.19 |

**S3 Table.**
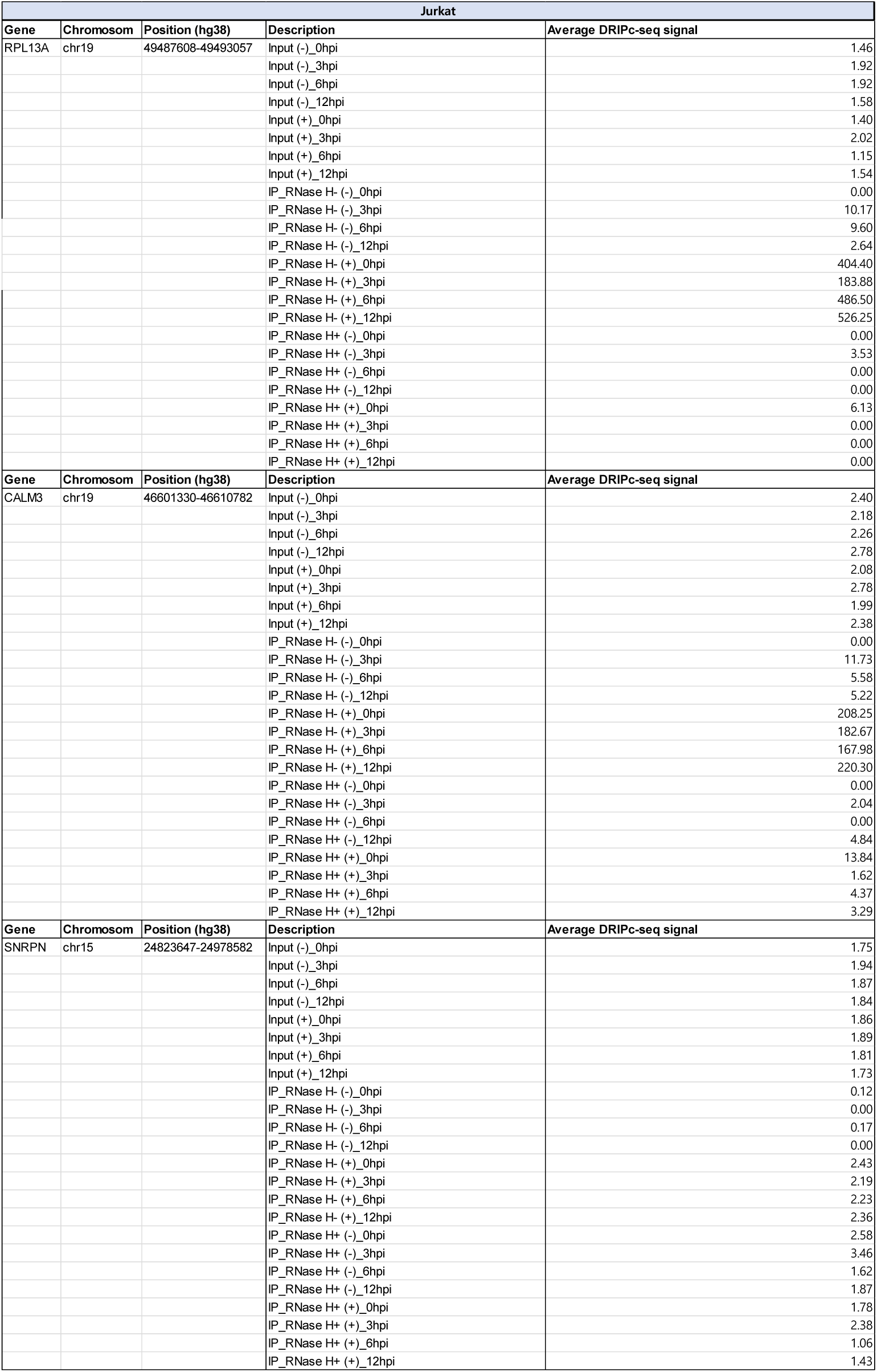
Chromosomal position and DRIPc-seq signal for referenced R-loop-positive and –negative regions in HIV-1 infected Jurkat cells.

**S4 Table.** RNA-seq analysis of relative gene expression levels of P1-3 and N1,2 R-loop regions.

|  | gene_symbol | 0hpi | 3hpi | 6hpi | 12hpi | 48hpi |
| --- | --- | --- | --- | --- | --- | --- |
| **P1** | TOR1AIP2 | 1 | 1.163832 | 1.247899 | 1.024926 | 0.619497 |
| **P2** | DVL1 | 1 | 0.781593 | 0.571348 | 0.901502 | 0.270459 |
| **P3** | PKN2 | 1 | 1.280974 | 1.891552 | 1.31842 | 1.515107 |
| **N1** | N/A | N/A | N/A | N/A | N/A | N/A |
| **N2** | CDK5RAP1 | 1 | 0.73977 | 0.775 | 1.143662 | 0.472377 |

**S5 Table.**
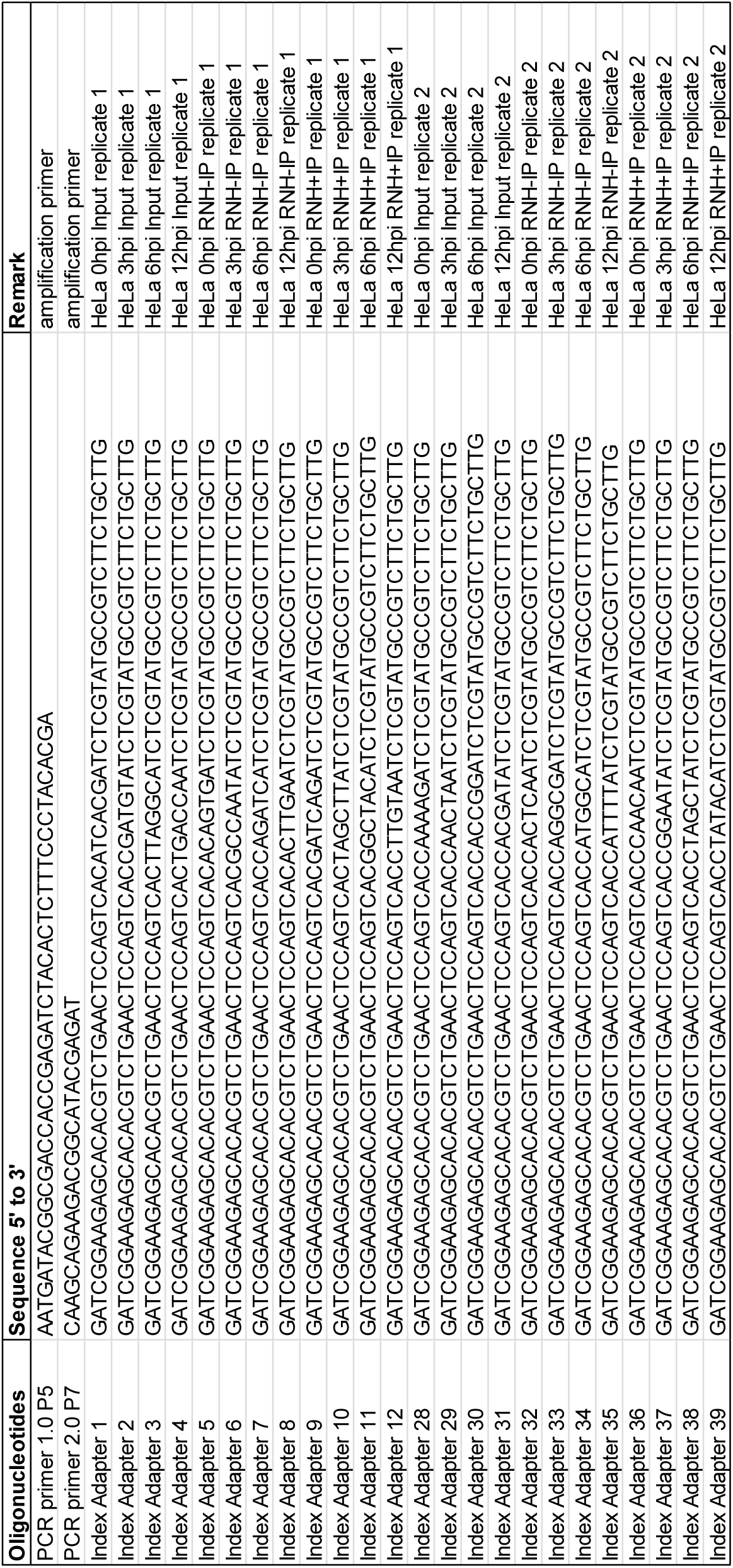
Oligonucleotides used for DRIPc-seq library construction.

**S6 Table.**
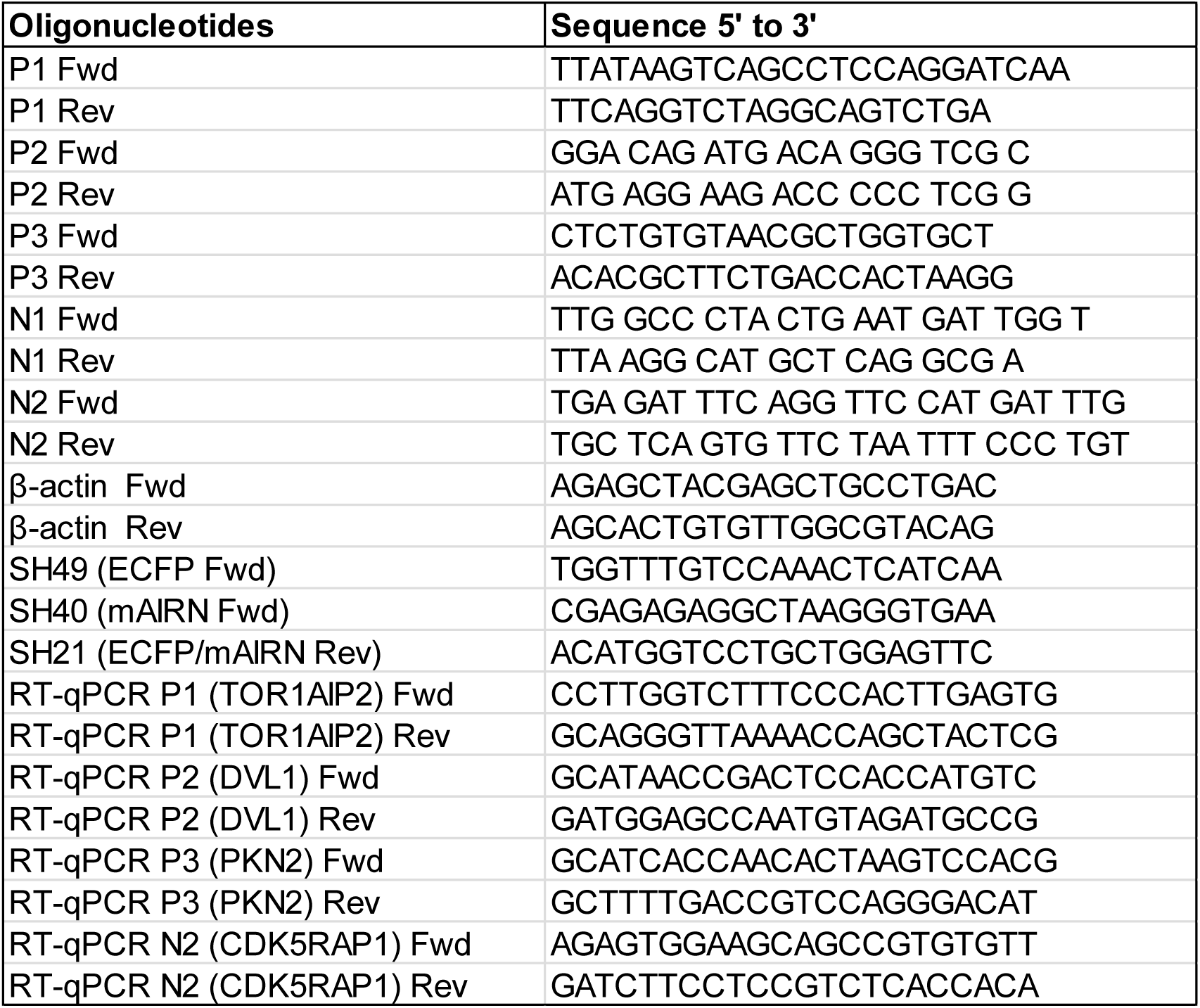
Primers used for qPCR.

**S7 Table.**
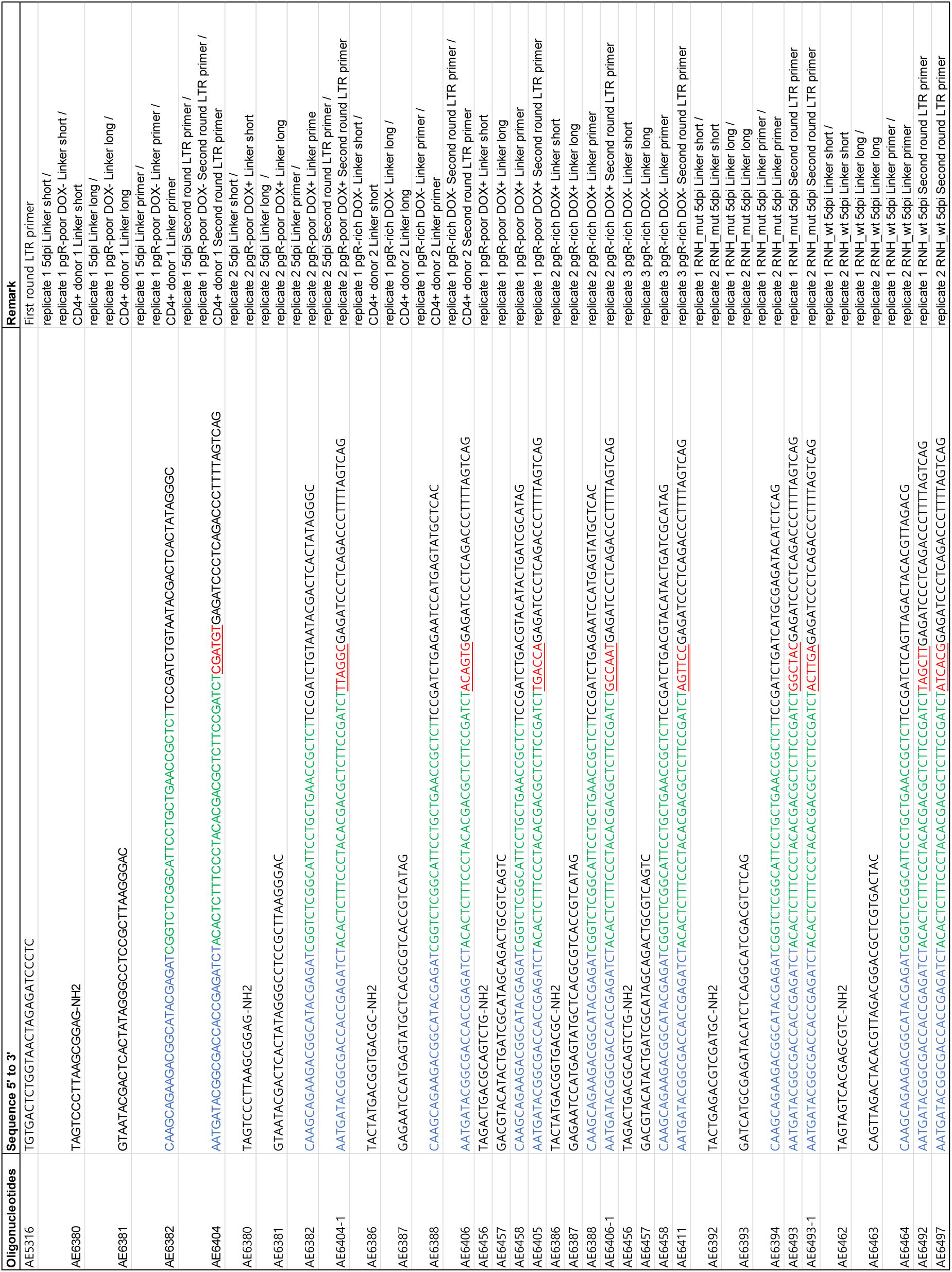
Oligonucleotides used for HIV-1 integration site sequencing library construct.

**S8 Table.**
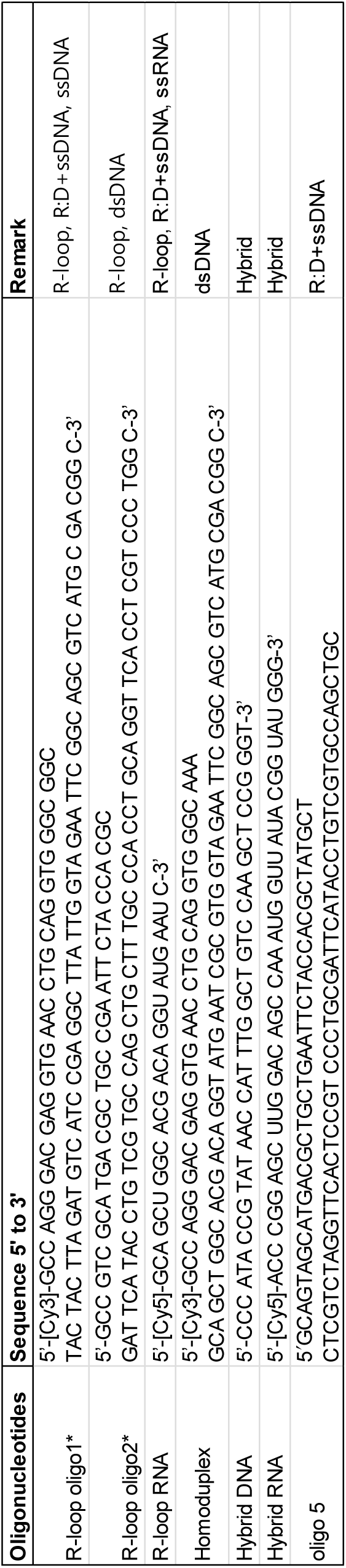
Oligonucleotides used for electrophoretic mobility shift assay substrate preparation.

**S9 Table.**
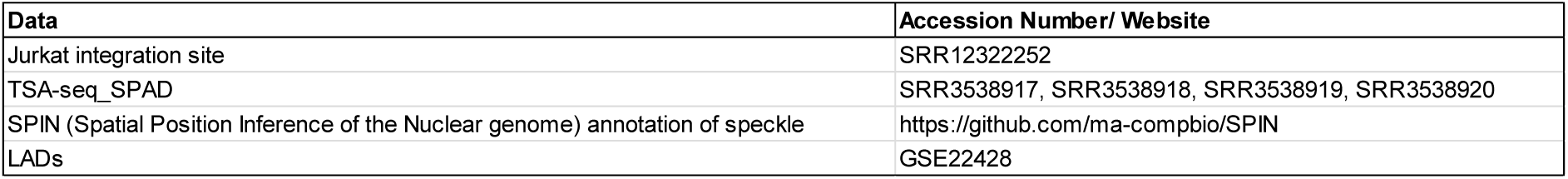
Accession numbers and data sources.

| Data | Accession Number/ Website |
| --- | --- |
| Jurkat integration site | SRR12322252 |
| TSA-seq_SPAD | SRR3538917, SRR3538918, SRR3538919, SRR3538920 |
| SPIN (Spatial Position Inference of the Nuclear genome) annotation of speckle | https://github.com/ma-compbio/SPIN |
| LADs | GSE22428 |

## References

1. W. E. Johnson, Origins and evolutionary consequences of ancient endogenous retroviruses. Nat Rev Microbiol 17, 355–370 (2019).

2. M. Lusic, R. F. Siliciano, Nuclear landscape of HIV-1 infection and integration. Nat Rev Microbiol 15, 69–82 (2017).

3. H. C. Chen, J. P. Martinez, E. Zorita, A. Meyerhans, G. J. Filion, Position effects influence HIV latency reversal. Nat Struct Mol Biol 24, 47–54 (2017).

4. K. B. Einkauf et al., Parallel analysis of transcription, integration, and sequence of single HIV-1 proviruses. Cell 185, 266–282 e215 (2022).

5. C. Jiang et al., Distinct viral reservoirs in individuals with spontaneous control of HIV-1. Nature 585, 261–267 (2020).

6. A. R. Schroder et al., HIV-1 integration in the human genome favors active genes and local hotspots. Cell 110, 521–529 (2002).

7. V. Achuthan et al., Capsid-CPSF6 Interaction Licenses Nuclear HIV-1 Trafficking to Sites of Viral DNA Integration. Cell Host Microbe 24, 392–404 e398 (2018).

8. A. Ciuffi et al., A role for LEDGF/p75 in targeting HIV DNA integration. Nat Med 11, 1287–1289 (2005).

9. G. A. Sowd et al., A critical role for alternative polyadenylation factor CPSF6 in targeting HIV-1 integration to transcriptionally active chromatin. Proc Natl Acad Sci U S A 113, E1054–1063 (2016).

10. B. Lucic et al., Spatially clustered loci with multiple enhancers are frequent targets of HIV-1 integration. Nat Commun 10, 4059 (2019).

11. B. Marini et al., Nuclear architecture dictates HIV-1 integration site selection. Nature 521, 227–231 (2015).

12. M. Kvaratskhelia, A. Sharma, R. C. Larue, E. Serrao, A. Engelman, Molecular mechanisms of retroviral integration site selection. Nucleic Acids Res 42, 10209–10225 (2014).

13. P. Cherepanov et al., HIV-1 integrase forms stable tetramers and associates with LEDGF/p75 protein in human cells. J Biol Chem 278, 372–381 (2003).

14. J. M. Coffin et al., Integration in oncogenes plays only a minor role in determining the in vivo distribution of HIV integration sites before or during suppressive antiretroviral therapy. PLoS Pathog 17, e1009141 (2021).

15. C. Niehrs, B. Luke, Regulatory R-loops as facilitators of gene expression and genome stability. Nat Rev Mol Cell Biol 21, 167–178 (2020).

16. E. Petermann, L. Lan, L. Zou, Sources, resolution and physiological relevance of R-loops and RNA-DNA hybrids. Nat Rev Mol Cell Biol 23, 521–540 (2022).

17. S. Hamperl, M. J. Bocek, J. C. Saldivar, T. Swigut, K. A. Cimprich, Transcription-Replication Conflict Orientation Modulates R-Loop Levels and Activates Distinct DNA Damage Responses. Cell 170, 774–786 e719 (2017).

18. P. A. Ginno, P. L. Lott, H. C. Christensen, I. Korf, F. Chedin, R-loop formation is a distinctive characteristic of unmethylated human CpG island promoters. Mol Cell 45, 814–825 (2012).

19. Y. W. Lim, L. A. Sanz, X. Xu, S. R. Hartono, F. Chedin, Genome-wide DNA hypomethylation and RNA:DNA hybrid accumulation in Aicardi-Goutieres syndrome. Elife 4, (2015).

20. R. Arora et al., RNaseH1 regulates TERRA-telomeric DNA hybrids and telomere maintenance in ALT tumour cells. Nat Commun 5, 5220 (2014).

21. T. Garcia-Muse, A. Aguilera, R Loops: From Physiological to Pathological Roles. Cell 179, 604–618 (2019).

22. L. A. Sanz et al., Prevalent, Dynamic, and Conserved R-Loop Structures Associate with Specific Epigenomic Signatures in Mammals. Mol Cell 63, 167–178 (2016).

23. F. Chedin, Nascent Connections: R-Loops and Chromatin Patterning. Trends Genet 32, 828–838 (2016).

24. C. Y. Lee et al., R-loop induced G-quadruplex in non-template promotes transcription by successive R-loop formation. Nat Commun 11, 3392 (2020).

25. H. O. Ajoge et al., G-Quadruplex DNA and Other Non-Canonical B-Form DNA Motifs Influence Productive and Latent HIV-1 Integration and Reactivation Potential. Viruses 14, (2022).

26. F. Chedin, C. J. Benham, Emerging roles for R-loop structures in the management of topological stress. J Biol Chem 295, 4684–4695 (2020).

27. I. K. Jozwik et al., B-to-A transition in target DNA during retroviral integration. Nucleic Acids Res 50, 8898–8918 (2022).

28. A. Ballandras-Colas et al., Multivalent interactions essential for lentiviral integrase function. Nat Commun 13, 2416 (2022).

29. L. A. Sanz, F. Chedin, High-resolution, strand-specific R-loop mapping via S9.6-based DNA-RNA immunoprecipitation and high-throughput sequencing. Nat Protoc 14, 1734–1755 (2019).

30. R. B. Jones et al., LINE-1 retrotransposable element DNA accumulates in HIV-1-infected cells. J Virol 87, 13307–13320 (2013).

31. S. Srinivasachar Badarinarayan et al., HIV-1 infection activates endogenous retroviral promoters regulating antiviral gene expression. Nucleic Acids Res 48, 10890–10908 (2020).

32. P. Lesbats, A. N. Engelman, P. Cherepanov, Retroviral DNA Integration. Chem Rev 116, 12730–12757 (2016).

33. A. Brussel, P. Sonigo, Analysis of early human immunodeficiency virus type 1 DNA synthesis by use of a new sensitive assay for quantifying integrated provirus. J Virol 77, 10119–10124 (2003).

34. A. Albanese, D. Arosio, M. Terreni, A. Cereseto, HIV-1 pre-integration complexes selectively target decondensed chromatin in the nuclear periphery. PLoS One 3, e2413 (2008).

35. A. Dharan, N. Bachmann, S. Talley, V. Zwikelmaier, E. M. Campbell, Nuclear pore blockade reveals that HIV-1 completes reverse transcription and uncoating in the nucleus. Nat Microbiol 5, 1088–1095 (2020).

36. P. A. Ginno, Y. W. Lim, P. L. Lott, I. Korf, F. Chedin, GC skew at the 5’ and 3’ ends of human genes links R-loop formation to epigenetic regulation and transcription termination. Genome Res 23, 1590–1600 (2013).

37. J. J. Kessl et al., HIV-1 Integrase Binds the Viral RNA Genome and Is Essential during Virion Morphogenesis. Cell 166, 1257–1268 e1212 (2016).

38. D. C. van Gent, Y. Elgersma, M. W. Bolk, C. Vink, R. H. Plasterk, DNA binding properties of the integrase proteins of human immunodeficiency viruses types 1 and 2. Nucleic Acids Res 19, 3821–3827 (1991).

39. A. Cristini, M. Groh, M. S. Kristiansen, N. Gromak, RNA/DNA Hybrid Interactome Identifies DXH9 as a Molecular Player in Transcriptional Termination and R-Loop-Associated DNA Damage. Cell Rep 23, 1891–1905 (2018).

40. T. Mosler et al., R-loop proximity proteomics identifies a role of DDX41 in transcription-associated genomic instability. Nat Commun 12, 7314 (2021).

41. R. Schrijvers et al., LEDGF/p75-independent HIV-1 replication demonstrates a role for HRP-2 and remains sensitive to inhibition by LEDGINs. PLoS Pathog 8, e1002558 (2012).

42. P. C. Stirling, P. Hieter, Canonical DNA Repair Pathways Influence R-Loop-Driven Genome Instability. J Mol Biol 429, 3132–3138 (2017).

43. M. L. Garcia-Rubio et al., The Fanconi Anemia Pathway Protects Genome Integrity from R-loops. PLoS Genet 11, e1005674 (2015).

44. M. Giannini et al., TDP-43 mutations link Amyotrophic Lateral Sclerosis with R-loop homeostasis and R loop-mediated DNA damage. PLoS Genet 16, e1009260 (2020).

45. S. Fu et al., HIV-1 exploits the Fanconi anemia pathway for viral DNA integration. Cell Rep 39, 110840 (2022).

46. D. Li, A. Lopez, C. Sandoval, R. Nichols Doyle, O. I. Fregoso, HIV Vpr Modulates the Host DNA Damage Response at Two Independent Steps to Damage DNA and Repress Double-Strand DNA Break Repair. mBio 11, (2020).

47. H. Bauby et al., HIV-1 Vpr Induces Widespread Transcriptomic Changes in CD4(+) T Cells Early Postinfection. mBio 12, e0136921 (2021).

48. K. Stopak, C. de Noronha, W. Yonemoto, W. C. Greene, HIV-1 Vif blocks the antiviral activity of APOBEC3G by impairing both its translation and intracellular stability. Mol Cell 12, 591–601 (2003).

49. D. Kmiec, F. Kirchhoff, Antiviral factors and their counteraction by HIV-1: many uncovered and more to be discovered. J Mol Cell Biol, (2024).

50. J. L. McCann et al., R-loop homeostasis and cancer mutagenesis promoted by the DNA cytosine deaminase APOBEC3B. 2021.2008.2030.458235 (2021).

51. S. A. Yukl et al., HIV latency in isolated patient CD4(+) T cells may be due to blocks in HIV transcriptional elongation, completion, and splicing. Sci Transl Med 10, (2018).

52. D. O. Passos et al., Cryo-EM structures and atomic model of the HIV-1 strand transfer complex intasome. Science 355, 89–92 (2017).

53. H. D. Nguyen et al., Functions of Replication Protein A as a Sensor of R Loops and a Regulator of RNaseH1. Mol Cell 65, 832–847 e834 (2017).

54. S. Andrews. (2010).

55. F. J. Felix Krueger, Phil Ewels, Ebrahim Afyounian, & Benjamin Schuster-Boeckler, FelixKrueger/TrimGalore: v0.6.7 – DOI via Zenodo (0.6.7). Zenodo. (2021).

56. M. Martin, Cutadapt removes adapter sequences from high-throughput sequencing reads. 2011 17, 3 %J EMBnet.journal (2011).

57. H. Li, R. Durbin, Fast and accurate short read alignment with Burrows-Wheeler transform. Bioinformatics 25, 1754–1760 (2009).

58. Y. Zhang et al., Model-based analysis of ChIP-Seq (MACS). Genome Biol 9, R137 (2008).

59. A. R. Quinlan, I. M. Hall, BEDTools: a flexible suite of utilities for comparing genomic features. Bioinformatics 26, 841–842 (2010).

60. W. Li et al., CPSF6-Dependent Targeting of Speckle-Associated Domains Distinguishes Primate from Nonprimate Lentiviral Integration. mBio 11, (2020).

61. H. M. Amemiya, A. Kundaje, A. P. Boyle, The ENCODE Blacklist: Identification of Problematic Regions of the Genome. Sci Rep 9, 9354 (2019).

62. A. Frankish et al., Gencode 2021. Nucleic Acids Res 49, D916–D923 (2021).

63. A. Dobin et al., STAR: ultrafast universal RNA-seq aligner. Bioinformatics 29, 15–21 (2013).

64. Y. Jin, O. H. Tam, E. Paniagua, M. Hammell, TEtranscripts: a package for including transposable elements in differential expression analysis of RNA-seq datasets. Bioinformatics 31, 3593–3599 (2015).

